# Visual information routes in the posterior dorsal and ventral face network studied with intracranial neurophysiology, and white matter tract endpoints

**DOI:** 10.1101/2020.05.22.102046

**Authors:** M Babo-Rebelo, A Puce, D Bullock, L Hugueville, F Pestilli, C Adam, K Lehongre, V Lambrecq, V Dinkelacker, N George

## Abstract

Occipito-temporal regions within the face network process perceptual and socio-emotional information, but the dynamics and information flow between different nodes of this network is still debated. Here, we analyzed intracerebral EEG from 11 epileptic patients viewing a stimulus sequence beginning with a neutral face with direct gaze. The gaze could avert or remain direct, while the emotion changed to fearful or happy. N200 field potential peak latencies indicated that face processing begins in inferior occipital cortex and proceeds anteroventrally to fusiform and inferior temporal cortices, in parallel. The superior temporal sulcus responded preferentially to gaze changes with augmented field potential amplitudes for averted versus direct gaze, and large effect sizes relative to other network regions. An overlap analysis of posterior white matter tractography endpoints (from 1066 healthy brains) relative to active intracerebral electrodes in the 11 patients showed likely involvement of both dorsal and ventral posterior white matter pathways. Overall, our data provide new insight on the timing of face and social cue processing in the occipito-temporal brain and anchor the superior temporal cortex in dynamic gaze processing.

## INTRODUCTION

Faces are critical social stimuli as they provide unique information about identity, emotional and mental states, and as such they are the primary focus of social attention. This social information is gleaned quickly, typically within a fraction of a second of seeing the face, as in a fleeting glance. There is a wealth of neuroimaging and neuropsychological research on the core and extended network for face processing (see Figure 4 of (Gobbini and Haxby 2007)), with information channeled along two main functional pathways, one based on identity and the other on the changeable aspects of faces such as gaze and emotional expression, consistent with the pioneering neuropsychological model of Bruce and Young (Bruce and Young 1986).

The core face network includes three regions: the inferior occipital gyrus, fusiform gyrus, and superior temporal sulcus (STS) (Haxby et al. 2000). The fusiform gyrus is thought to extract the non-variant aspects of facial features and their spatial relations inherently associated with an individual’s identity. Human functional magnetic resonance imaging (fMRI) studies have identified a series of three face sensitive patches on human fusiform gyrus (Pinsk et al. 2009; Engell and McCarthy 2013; Grill-Spector et al. 2017). Meta-analyses indicate that acquired prosopagnosia in humans typically involves the posterior fusiform and inferior occipital gyri (Bouvier and Engel 2006). The core face network also includes the STS, a region associated with multisensory integration (Beauchamp et al. 2004), the processing of biological motion (Bonda et al. 1996), facial motion (Puce et al. 1998), and social attention (Allison et al. 2000). Within the framework of the core face network, the STS deals with the dynamic aspects of the human face - such as gaze changes and emotional expressions (Gobbini and Haxby 2007). According to one putative organization scheme, the inferior occipital and fusiform gyri are proposed to lie within the ventral visual stream, whereas the STS is a dorsal visual stream structure (Bernstein and Yovel 2015). It has also been speculated that the inferior occipital gyrus could be a potential entry point to the system (Haxby et al. 2000; Fairhall and Ishai 2007) feeding both the fusiform gyrus and STS. However, this hierarchical view is put into question by evidence from neuropsychological lesion studies and information within the core face network may be bridged between the ventral and dorsal visual streams bypassing the inferior occipital region (Rossion, 2008; Atkinson and Adolphs, 2011; Weiner et al., 2016). A very recent framework has added a third visual pathway to the scheme—where information from V1 makes its way via MT/V5 to the STS—based on multimodal data from non-human primates (Pitcher and Ungerleider, 2021).

The time course and nature of the interactions within the core face network remain unknown (e.g., Kennedy and Adolphs 2012; Stanley and Adolphs 2013) and there are some potentially conflicting models that are based heavily on fMRI data (e.g. Duchaine & Yovel 2015; Jiang et al., 2011). Intracerebral field potential studies (iEEG) can provide unique information in that domain. Most of the iEEG studies of the face network have focused on the ventral occipitotemporal cortex (predominantly fusiform gyrus), and to a lesser extent on the lateral occipitotemporal cortex (for recent examples, see Li et al. 2019; Rangarajan et al. 2020; Schrouff et al. 2020). Previous studies have documented a series of local field potential components including a face-sensitive N200 in response to isolated static faces (e.g., Allison et al. 1994; Halgren et al. 1994; Puce et al. 1999; Barbeau et al. 2008; Pourtois et al. 2010). The N200 has been shown to be equivalent to the scalp EEG N170 in response to upright faces in the same individuals (Rosburg et al. 2010). It is relatively invariant to habituation, priming and other top-down factors relative to later face-selective field potentials, which occur at latencies as late as 700 ms post-stimulus (Puce et al. 1999). This co-occurrence of early and late neurophysiological activity at the *same cortical locations* is likely to produce complex fMRI signals—potentially confounding the construction of models of cortical information flow that rely on fMRI data.

In real life, face processing involves the simultaneous processing of dynamic, short duration social cues, such as emotional expressions and social attention (direction of gaze). This represents an important experimental challenge in the laboratory. Our previous EEG and MEG studies have used continuously presented faces that change dynamically (Puce et al. 2000; Conty et al. 2007; Ulloa et al. 2014; Huijgen et al. 2015; Latinus et al. 2015; Rossi et al. 2015). We reported increased scalp N170 ERP and M170 MEG responses to viewing gaze aversion relative to direct gaze when the head faces the observer with a dynamic paradigm (Puce et al. 2000; Ulloa et al. 2014; Latinus et al. 2015). Interestingly, there is a dearth of iEEG studies examining lateral temporal cortex, including the posterior STS—an essential part of the face network, particularly when it comes to dynamic social cues. To our knowledge, Caruana et al. is the only group that have reported iEEG field potentials to a dynamic gaze change task from the STS, observing larger field potentials at around 200 ms post-stimulus to viewing averted gaze relative to direct gaze (Caruana et al. 2014). Thus, from the neurophysiological perspective, there is a lack of knowledge regarding the unfolding in time of the processing of faces and dynamic social facial cues (i.e. gaze and emotion) throughout the core face processing network. In particular, the exact role of the superior temporal sulcus (STS) in the coding of social information and how this relates to the timing of activity in other core face-network structures is poorly understood.

In addition to a limited understanding of the functional interactions between the structures of the face network, the structural interconnections across this network have not been well documented, although this has attracted growing interest recently (Thomas et al. 2009; Grill-Spector et al. 2017; Wang and Olson 2018). Direct and indirect white matter connections within the core face network exist between the occipital and fusiform face responsive regions (inferior occipital gyrus/occipital face area or OFA, posterior fusiform/fusiform face area or FFA, and mid-fusiform gyrus) via the inferior longitudinal fasciculus and shorter range occipito-temporal tracts (Catani et al., 2003; Pyles et al. 2013; Grill-Spector et al. 2017). Additionally, vertical white matter tracts, including the posterior aspect of the arcuate fasciculus and anterior portions of the vertical occipital fasciculus, might play a role in linking visually-responsive structures of the dorsal and ventral pathway (Weiner et al. 2016, 2017). Some areas within the face network show direct connections with structures outside the network. For example, there is a direct white matter connection between the posterior fusiform gyrus and the intraparietal sulcus via the vertical occipital fasciculus (Yeatman et al. 2014; Takemura et al. 2017; Weiner et al., 2016; Grill-Spector et al. 2017). Yet, the STS has no known direct connections to the fusiform gyrus (Ethofer et al. 2011; Gschwind et al. 2012; Pyles et al. 2013; Grill-Spector et al. 2017), leaving open the question of how interactions take place between the core face processing regions. These interactions may, in part, proceed through the extended face network, which includes structures such as posterior parietal cortex and temporo-parietal junction, the amygdala, the insula, and anterior temporal cortex. Furthermore, as already noted, a third visual pathway could bring information from V1 to the STS (Pitcher and Ungerleider, 2021). It has been suggested that information flow in the face network via short range white matter tracts is also important (Wang et al. 2020). All in all, there remains a need to consider the role of white matter tract connections in information flow through the face network. This is particularly pertinent for neurophysiological studies where subtle latency differences across regions and stimulus conditions could arise from information flow through different routes in the face connectome.

In this study, we attempted to fill the knowledge gap on function and connectivity of the face network, by analyzing a large iEEG dataset, of which only the amygdala data had been investigated so far (Huijgen et al. 2015). Specifically, we aimed at addressing the following questions:

i) Where are the predominant sites that respond to face onset and changes in gaze and emotion across the face network? How does waveform morphology, amplitude and latency alter as a function of these facial attributes?
ii) What parts of the face network are sensitive to changes in gaze direction vs. emotional expression?
iii) What are the likely routes of information flow across structures in the face network?

We expressed the active contacts of our iEEG dataset in a bipolar configuration, adopting the inferior occipital cortex, fusiform cortex, superior temporal cortex and inferior temporal cortex as our four regions of interest (ROIs). The inferior temporal cortex is more rarely considered, but lies between fusiform and superior temporal cortices and is also responsive to faces (Ishai et al. 2005). Given the high-signal-to-noise ratio of N200, based on previous literature, we focused on N200 amplitude and latency as metrics to study sensitivity and timing differences within the core face processing system. Then, for assessing likely routes of information flow in the face network, we complemented our iEEG dataset with data from 1066 healthy subjects of the Human Connectome Project (HCP; Van Essen et al. 2013). We calculated common posterior brain white matter tract pathway endpoints in MNI coordinate space (see Bullock et al. 2019) from the HCP subjects and then superimposed tract pathway endpoints with the relative locations of active bipolar sites from our patient sample.

## MATERIALS AND METHODS

### Patients

Eighteen patients with drug-refractory epilepsy were originally included in this study. These patients were implanted with depth electrodes, as part of their clinical pre-surgical evaluation at the Epileptology Unit in the Pitié-Salpêtrière Hospital, Paris, France. Implantation sites were only based on clinical criteria. All patients provided written informed consent to take part in the experiment. The study was approved by the local ethics committee (CPP Ile-de-France VI) and adheres to the principles of the Declaration of Helsinki.

We excluded 7 of the 18 patients from the analysis because of either high levels of interictal epileptic activity leading to an insufficient number of trials per condition (6 patients), or grossly disrupted brain anatomy due to the presence of a large lesion (1 patient). The remaining 11 patients (mean age = 30 years, range 20-48; five females; see Table 1) were included in the analyses. All had normal or corrected-to-normal vision. Patients 1 to 17 were tested between 2009 and 2012 and were in the cohort described by Huijgen and colleagues in 2015, from which the amygdala data of 5 patients were analyzed (Huijgen et al. 2015). Patient 18 was tested in 2018. Unlike in 2015, here, we performed an extensive analysis of the intracerebral recordings from sites spread throughout the ventral and lateral occipitotemporal cortices.

**Table 1.** Clinical characteristics of the 11 patients included in data analysis. N AED=number of anti-epileptic drugs. STAI=State-Trait Anxiety Inventory scores; M=male; F=female; L=left; R=right; post=posterior.

| Patient | Sex | Age (years) | Handedness | Seizure onset age (years) | Medication (N AED) | Epileptogenic zone | STAI |
| --- | --- | --- | --- | --- | --- | --- | --- |
| 1 | M | 25 | L | 10 | 3 | R post temporal | 36 |
| 2 | F | 24 | R | 5 | 2 | R temporo-occipital | 32 |
| 3 | F | 22 | R | 8 | 3 | R basal temporal | 29 |
| 8 | F | 28 | R | 24 | 2 | R temporo-polar & amygdalo-hippocampal | 40 |
| 10 | M | 25 | R | 15 | 4 | R temporo-occipital | - |
| 12 | M | 20 | R | 16 | 2 | L basal temporal | 21 |
| 14 | M | 48 | R | 22 | 2 | R temporo-lateral & multi- focal (R orbital, opercular) | 38 |
| 15 | M | 37 | L | 1 | 3 | R post & lateral occipital | 26 |
| 16 | F | 44 | R | 19 | 2 | R basal temporal | 35 |
| 17 | M | 31 | R | 12 | 3 | R post superior temporal | 26 |
| 18 | F | 31 | R | 26 | 2 | L temporal periventricular | 28 |

### Stimuli and experimental protocol

The experimental paradigm (Fig. 1) has been previously described in detail, as it has been used with magnetoencephalography in healthy subjects (Lachat et al. 2012), and in our amygdala iEEG recordings in epileptic patients (Huijgen et al. 2015). Sixteen different unfamiliar greyscale faces served as stimuli in a Posner-like task design consisting of a series of visual stimulus transitions. Each face stimulus had exemplars with direct and averted gaze, combined with a happy, fearful, or neutral expression. The neutral face with direct gaze exemplar always served as the initial face stimulus (Face 1) and was followed by the same face with either a happy or fearful expression and with averted or direct gaze (Face 2). This stimulus transition produced an apparent motion (and emotion) effect. Experimental trials were generated so that gaze changes were equalized across emotions i.e. equal numbers of gaze changes to the left and right were presented within both emotion conditions (see (Huijgen et al. 2015)).

**Figure 1.**
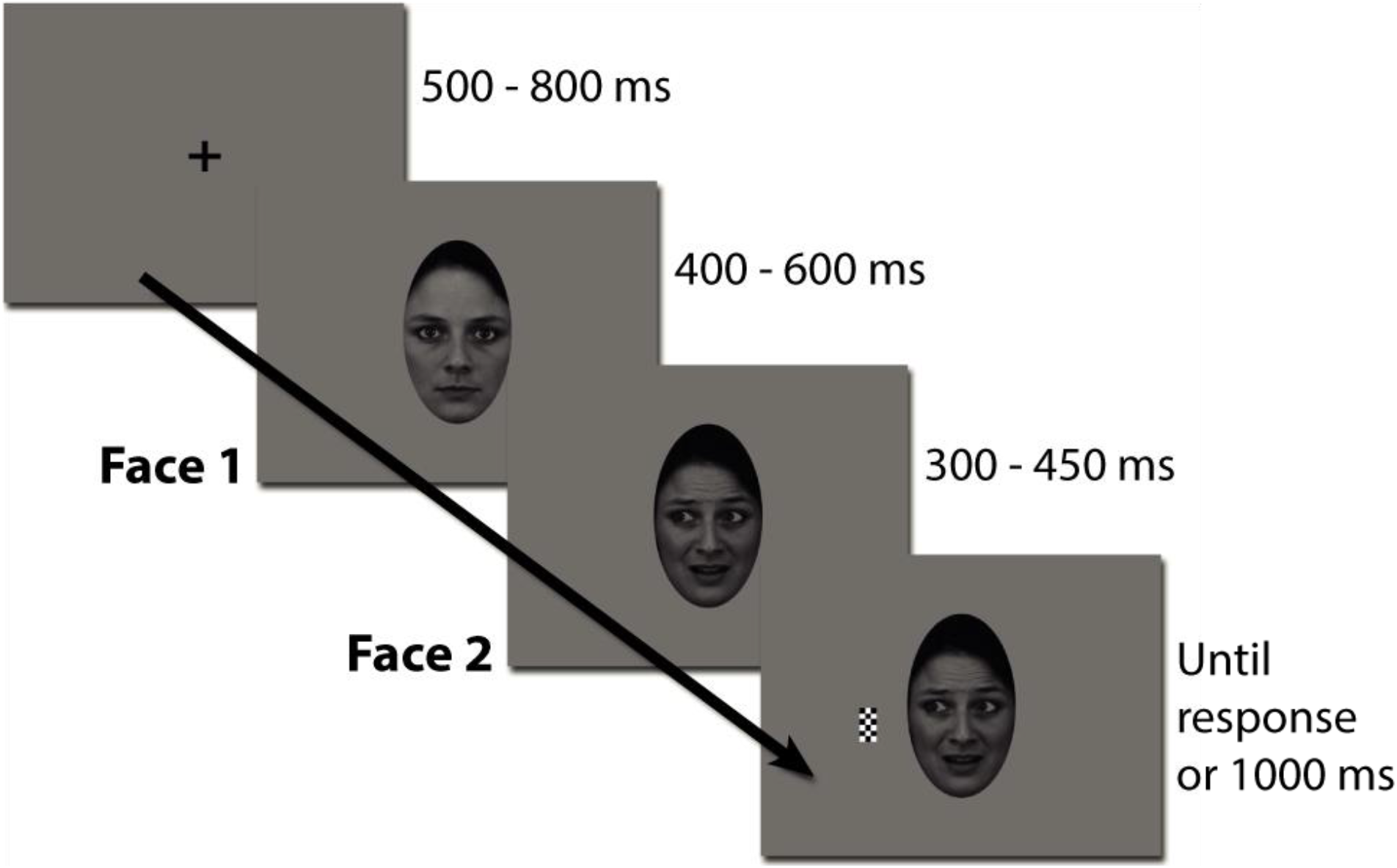
Experimental paradigm. A trial began with a fixation cross that was replaced by a neutral face with direct gaze (Face 1). After a variable delay, the face turned into happy or fearful with or without gaze aversion (Face 2), in an apparent motion manipulation. In 89% of the trials, a checkerboard would then appear on the left or right of the face. The patient had to press a button as quickly as possible after the appearance of the target checkerboard.

Each trial began with a black central fixation cross presented on a grey background for a random duration between 500 and 800 ms, followed by the onset of a neutral face looking directly at the observer (Face 1). After a variable delay of 400-600 ms duration (except for Patient 1 in whom the delay was fixed at 500 ms), the face became happy or fearful, with or without an associated gaze direction change (Face 2). Then, in 89% of the trials, after a randomly chosen interval of 300, 350, 400, or 450 ms, a checkerboard target appeared either on the right or on the left side of the face, that is, either on the side to which the eyes looked (congruent target condition) or on the opposite side (incongruent target condition). In the remaining 11% of the trials, the target was omitted (catch trials). The task was a mere detection task: Subjects had to press a response button as quickly as possible to the appearance of the target (whatever its location) and to refrain from pressing the button when no target was present. The stimulus display remained on screen until the subject’s response or a maximum of 1s had elapsed. Fixation had to be maintained throughout the entire trial and the subjects were asked to refrain from blinking during stimulus presentation. Subjects were required to perform up to eight blocks of this task. Each block comprised 108 trials (54 happy/fearful, 36 left/right/direct gaze, including 12 catch trials) with randomized order of the stimulus condition presentation (total of 864 trials, including 96 catch trials). Patients were given an option to rest between successive blocks.

The task was presented using either a cathode ray tube screen (Patients 1-17, resolution=1024×768) placed at 60 cm distance from the patient or a laptop screen (Patient 18, resolution=1920×1200) at 56 cm viewing distance. All patients viewed the centrally presented face stimuli with a visual angle of 4.3° (horizontal)× 7.3° (vertical). The target stimulus subtended a visual angle of 0.2°× 0.4° and was presented at a visual angle of 6.5° from the central fixation cross. The delay of stimulus presentation introduced by the laptop screen was constant at 27 ms and was corrected offline.

After completing the task, patients proceeded to fill out the State-Trait Anxiety Inventory (STAI Form Y, Self-Evaluation Questionnaire (Spielberger et al. 1983; Ansseau 1997)), which revealed that their anxiety scores were within the normal range (see Table 1; mean ± SEM = 31.1 ± 1.9).

### Behavioral data

Our main aim for measuring behavior was to ensure that the patients were performing the task correctly and attending to the stimuli. We computed the hit rate as the proportion of correct target detections and the false alarm rate as the proportion of button presses on catch trials. We excluded the responses that occurred prior to target presentation (range across patients: 0-5). For reaction time (RT) to targets, we discarded trials where the RT was above or below 3 standard deviations of the subject’s mean RT. This analysis showed that the 11 patients were able to easily detect the target: they made 99% (SEM = 0.6) hits and 3% (SEM = 1) false alarms (see Supplementary Table 1). The mean (± SEM) reaction time for target detection across patients was 357 ± 20 ms.

### Recordings

Patients were implanted with stereotactic depth electrodes, each containing 4 to 12 recording sites (AD-Tech, electrode width: 1 mm, inter-contact spacing: 10 mm for Patients 1, 2, 3, 8, 12 or 5 mm for Patients 10, 14, 15, 16, 17, 18) (Mathon et al. 2015). Intracerebral EEG (iEEG) data were acquired for Patients 1-3 with Nicolet 6000 system (Nicolet-Viasys, Madison, WI, USA) at a sampling rate of 400 Hz (bandpass: 0.05-150 Hz). The remaining patients were recorded using Micromed System 3 Plus (SD LTM 64 BS, Micromed S.p.A., Italy), at a sampling rate of 1024 Hz (bandpass: 0.15-250 Hz). The reference electrode was located between Fz and Cz on the scalp and a ground electrode was placed on the chest. Selected 10-10 scalp electrode locations also formed part of the recording montage in each patient, as allowed by their intracranial implantation. Electro-oculographic electrodes were also added to the recordings for Patients 2, 10, and 17.

Henceforth in this manuscript, we refer to ‘*electrodes’* as the implanted depth electrodes themselves, to ‘*contacts’* as the monopolar recording sites located along the electrodes, and to (bipolar) *’sites’* as the virtual recording site resulting from bipolar derivation of the monopolar recordings (see below, *Data preprocessing: Filtering, artifact identification and rejection, epoching and z-scoring*).

### Intracerebral electrode localization

Post-implantation high-resolution T1-weighted MRI scans (at 1.5 Tesla) were obtained for each patient. In-plane voxel sizes were 0.938 mm× 0.938 mm. Slice thickness was 0.7 mm in Patients 1 and 17, 1.2 mm in Patient 15, 1.3 mm in patients 8, 12, 14 and 16, and 1.5 mm in Patients 2, 3 and 10. Patient 18 had a 2 mm iso-voxel scan.

These T1 MRI scans were normalized using SPM 12 (Wellcome Centre for Human Neuroimaging, UCL, London) running under MATLAB 2015b (The Mathworks, Inc., Natick, MA, USA). The first step of analysis was intracerebral contact localization *prior to undertaking any analysis of the electrophysiological data*. For this, two observers (AP and MBR) visually labelled each recording contact in terms of sulci, gyri, and white matter, by visual inspection of the individual patient’s anatomy (based on the original post-implantation MRI), using the Duvernoy’s atlas of the human brain (Duvernoy 1992). They then determined the MNI coordinates for each electrode recording contact on the normalized post-implantation MRIs. Electrode contacts that were outside of the brain were identified, so that they could be excluded from further analyses.

Our analyses (as detailed below) focused on bipolar neurophysiological derivations. The MNI coordinates for bipolar sites were defined as the *mid-point* between the two corresponding monopolar contacts (see Supplementary Methods S1.1). Bipolar sites were then grouped according to four anatomical regions of interest (ROIs) for subsequent electrophysiological data analyses, including the three regions of the core face processing network (fusiform cortex, inferior occipital cortex and superior temporal cortex; (Haxby et al. 2000; Gobbini and Haxby 2007)) and an additional neighboring region (inferior temporal cortex). More precisely, each region included the following set of structures:

– ***Fusiform cortex (FC)*** included all bipolar sites labelled as fusiform gyrus, occipitotemporal sulcus, collateral sulcus, and mid-fusiform sulcus;
– ***Inferior occipital cortex (IOC)*** included the inferior occipital gyrus, inferior occipital sulcus, lateral occipital sulcus, mid-occipital gyrus, fourth occipital gyrus, and transverse posterior collateral sulcus;
– ***Superior temporal cortex (STC)*** included the superior temporal sulcus, middle temporal gyrus, and superior temporal gyrus;
– ***Inferior temporal cortex (ITC)*** included the inferior temporal sulcus and inferior temporal gyrus.

The toolbox iso2mesh (http://iso2mesh.sourceforge.net; (Fang and Boas 2009)) running under Matlab 2015b (The Mathworks, Inc., Natick, MA, USA) was used to illustrate electrode localization in all 3D brain views. Figure 2 shows the distribution of bipolar sites in terms of which patient they belong to (Fig. 2A), and in which ROI (Fig. 2B, 2C) they are located. See also Supplementary Table 2 for the number of bipolar sites that were analyzed in each region of interest for each patient.

**Figure 2.**
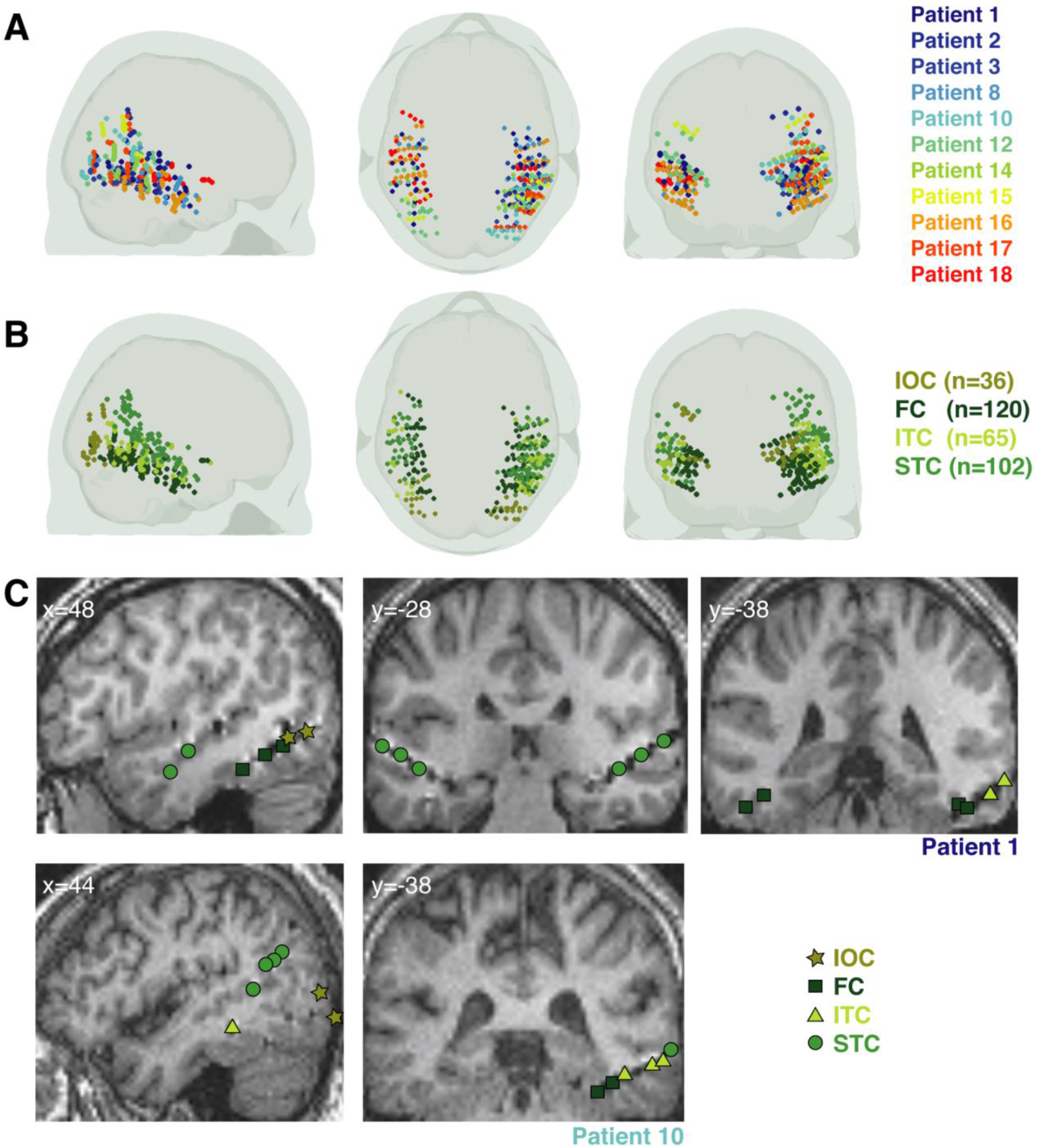
Occipitotemporal cortex sampling across the 11 patients. **A. Bipolar sites across patients**. The sagittal, axial and coronal views of the brain illustrate the coverage obtained with the 323 bipolar sites retained for analysis across the 11 patients. The sites are color-coded as a function of the patient to which they belong. **B. Bipolar sites across ROIs.** The same sites as in **A** are depicted, except that here they are color-coded as a function of the ROI in which they were localized: IOC, Inferior Occipital Cortex; FC, Fusiform Cortex; ITC, Inferior Temporal Cortex; STC, Superior Temporal Cortex. The number (n) of bipolar sites comprised in each ROI is indicated in parentheses. **C. Single patient illustration of bipolar site localization.** These (normalized) MRI images show some of the bipolar sites responding to faces for Patient 1 and Patient 10. For visualization purposes, these have been projected to the same x or y coordinate. The color and shape code the corresponding ROI.

Additionally, we also examined data from intracerebral electrodes located in the intraparietal sulcus region (IPS)—a region known to respond to dynamic face parts (Puce et al. 1998). However, since the number of sites was very small in this region (see Supplementary Table 2), we restricted our analysis in this region and present it in Supplementary Materials (See Supplementary Results S2.4).

### Data preprocessing: Filtering, artifact identification and rejection, epoching and z-scoring

All preprocessing and analyses of intracerebral EEG data were performed using Matlab 2015b and the FieldTrip toolbox ((Oostenveld et al. 2011), version 20180129), running in the Linux environment, unless specified otherwise.

Initial data review was performed on monopolar recordings. First, recording contacts that were consistently noisy (exhibiting frequent interictal epileptic activity or artifacts), were discarded from further analysis. Data were filtered with a high-pass cut-off of 0.3 Hz (order 4 Butterworth filter). We also applied two Notch filters (48-52 Hz and 58-66 Hz, Butterworth filters with order of 4) to exclude any remaining electrical noise.

To facilitate artifact detection, monopolar data were epoched from 400 ms before fixation to 1s after target onset, resulting in a single long epoch per trial. Epochs with signal amplitudes exceeding a voltage threshold of ± 750 μV were automatically excluded from further analysis. The remaining epochs were visually inspected, and abnormal activity (epileptic or muscle activity, electrical artifacts) was further identified. Epochs containing an artifact between 500 ms before Face 1 onset and 300 ms after target onset were discarded. Data from all contacts were visualized simultaneously, to more effectively exclude contacts with persistent interictal epileptic activity and to detect possible propagation of epileptic activity to contacts where interictal spikes might not necessarily be clearly visible. Trials with suspected propagation were excluded from further analysis. Blink detection was performed on the scalp electrode showing the clearest blink signal, using the semi-automatic interactive procedure implemented in FieldTrip (*ft_artifact_zvalue*). To sum up, trials were excluded based on the following criteria: 1) ± 750 μV threshold crossing; 2) presence of epileptic iEEG activity and other artifacts between 500 ms before Face 1 onset and 300 ms after target onset; 3) presence of blinks in this time window.

In addition, we excluded trials with aberrant behavioral responses (misses, false alarms, responses before target onset, and RTs above or below 3 standard deviations of the subject’s mean RT). Any block with less than 50% remaining trials was excluded from the analysis. The resulting number of retained trials per patient is presented in Supplementary Table 1.

We then applied an additional low-pass filter with cut-off of 40 Hz (order 6 Butterworth filter). Data that were originally acquired at 1024 Hz were downsampled to 400 Hz, to equate temporal resolution of data across all patients. All recording contacts were then re-expressed as bipolar derivations by subtracting the signal of two consecutive monopolar contacts (deeper-shallower, on the same depth electrode shaft). This procedure minimizes influences of volume-conduction and emphasizes local signals (Lachaux et al. 2003).

Finally, we extracted the bipolar data elicited to Face 1 and Face 2, respectively, by subdividing the initial long epoch. For this, we extracted electrophysiological data from 100 ms pre-stimulus to 400 ms post-stimulus for each considered stimulus onset in each trial. The data were z-scored relative to baseline (defined as the 100 ms preceding each considered stimulus onset, trial by trial).

### Analysis of responsiveness to Face 1 and Face 2

To identify the sites that were responsive to Face 1 and/or Face 2, we used a two-step procedure. First, we identified the sites that showed a minimal response to either Face 1 or Face 2 (or both), defined as an activity level averaged across trials exceeding a liberal threshold of Z= ±1 (i.e. 1 standard deviation relative to the baseline) in the time window of 0 to 400 ms relative to the relevant face stimulus onset. Second, we tested if activity at these sites significantly differed from zero. For this, we performed a cluster-based permutation t-test against zero across time at each site (Maris and Oostenveld 2007; see Supplementary Methods S1.2 for details). This procedure intrinsically corrects for multiple comparisons over time. Moreover, to account for the number of sites tested in each region of interest (ROI), we applied an additional Bonferroni correction: we considered the Monte-Carlo p-value as statistically significant if it was inferior to 0.05/N, where N was the total number of sites tested in the right or left ROI to which the tested site belonged. The sites showing at least one cluster of activity statistically different from zero between 0 and 400 ms after Face 1 and/or Face 2 were considered as *responsive* to the corresponding face stimulus.

We then tested for differences in proportions of responsive sites observed in each ROI. These results are presented in Supplementary Results S2.3.

### Analysis of Event-Related Potentials (ERPs) in response to Face 1 and Face 2

The z-scored, bipolar local field potential signals obtained at each responsive site were averaged across trials in response to Face 1 and Face 2, respectively.

#### ERP waveform morphology

We examined ERP morphology along the extended posterior-to-anterior span of the ITC, FC, and STC in the right hemisphere. For this, we analyzed ERPs in each ROI by grouping them according to their y-coordinates, into 5 slices of 10 mm from y=−81 to y=−31 (MNI coordinates) and a sixth slice containing the remaining more anterior sites (that is, with y>−31). For IOC, the y-range was limited and all sites within this ROI were grouped together.

To visualize the overall ERP waveforms in each slice of the ITC, FC and STC, and in the IOC, we first standardized the waveforms according to their polarity. For this, in each above-defined slice of the ITC, FC, and STC, and in the IOC, we computed an initial mean ERP across all site of the considered region, averaging absolute amplitude values. We picked the latency of the maximal activity within the broad time window of expected major peaks of activity (0 to 170 ms for Face 1 and 2 responses in IOC and for Face 1 responses in FC slices; 210 to 400 ms for Face 1 responses in STC slices; 0 to 210 ms for Face 2 in FC and STC slices and for Face 1 and 2 in ITC slices). Then, we computed the average ERP amplitude within ± 25ms around this latency, at each site of the considered region. If this average amplitude was positive, we multiplied the whole ERP timecourse by −1; if it was negative, it was kept unchanged. This allowed us to visualize the rectified ERP waveform at each individual site before computing the final average ERP in each region (Fig. 4). This was necessary because adjacent sites show similar waveform but of opposite polarity (i.e. sign) when they are located either side of a local generator (or dipole). While such polarity reversals are important because they are indicative of the presence of a local neural generator, they prevent assessment of the overall ERP waveform morphology across sites in regions of interest. Our rectification procedure allowed circumventing this issue.

#### ERP amplitude analysis

The amplitude of the early negative ERP response to Face 1 and Face 2 was analyzed at the sites where ERP peaks could be clearly identified in each ROI of the right hemisphere, that is: in slices A to E (y=−81 to −32 mm) for the FC, in slices B to D (y=−71 to −42 mm) for ITC, in slices C to E (y=−61 to −32 mm) for STC, and on all sites for IOC (see Fig. 4). In each included slice and in IOC, we extracted the peak latency of the early (negative) ERP in response to Face 1 and Face 2 respectively, from the ERP time course averaged across the sites of the considered region, as described above. We then measured the mean amplitude ± 25ms around this peak latency on a trial-by-trial basis, at each site (and without any rectification). Effect size for the amplitude in each condition of interest (here Face 1 and Face 2) for each site was then computed in the form of Cohen’s d using the following formula:

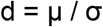

where μ is the average of the activity across trials and σ the standard deviation.

As the sign of d is dependent on the arbitrary polarity of the signal, we considered the absolute values of d. These were averaged across the sites considered within each ROI. We interpreted the magnitude of Cohen’s d according to Sawilowsky (2009), where a value of d=0.2 is defined as small, d=0.5 is medium and d=0.8 is large.

#### ERP latency analysis

Because peak latencies can be difficult to determine at the single site level, we used the jackknife procedure described in Miller et al (Miller et al. 1998) to estimate and compare latencies of the ERPs obtained for each experimental condition, in each ROI (see (Lochy et al. 2018) for a similar approach). We considered the rectified ERP time course averaged across the included sites in each ROI and measured the latency of the maximum negative ERP peak for Face 1 and for Face 2 on these time courses. The difference in the peak latency between Face 1 and 2 conditions was computed. The jackknife approach consists in repeating this procedure but for the average of all sites minus one: we iterated the procedure n times, by successively leaving out each of the n sites comprised in the considered ROI. This allowed us to obtain n latencies and to compute the corresponding standard deviation, for each condition. Then, to statistically compare the latencies between conditions, we derived the following t-value (Miller et al. 1998):

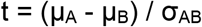

where μA − μB is the difference in mean peak latency between conditions A and B (here, Face 1 and Face 2), and σ_AB_ is the jackknife standard deviation computed as follows:

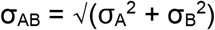

The degrees of freedom are equal to the total number of sites considered for conditions A and B minus 2. The 95% confidence interval and the two-sided p-value were then derived from the t-value and σ_AB_ to assess statistical significance.

The same procedure was applied to compare latencies *between ROIs* taken 2-by-2 for each experimental condition (Face 1 and Face 2). The p-values were then Bonferroni-corrected for multiple comparisons corresponding to the number of tests performed (that is, the threshold for significance was set to 0.05/n, with n=6) for each experimental condition. In addition, we also computed an ANOVA across the 4 ROIs for each Face condition, as described in (Ulrich and Miller, 2001).

### Analysis of the effects of emotion and gaze on ERPs

To analyze the modulation of ERPs to Face 2 as a function of emotion and gaze conditions, we averaged the z-scored bipolar EEG data in response to Face 2 separately for the fearful and happy faces with averted and direct gaze.

First, to test for the effects of emotion and gaze, we performed a 2-by-2 ANOVA across trials at each time point between 0 and 400 ms, on each site identified as responsive to Face 2 in the preceding analyses. This ANOVA included emotion (happy, fear) and gaze (direct, averted) as between-trial factors. It was implemented with the same clustering procedure as used above to correct for multiple comparisons over time.

The proportion of sites where statistically significant effects of emotion and/or gaze were observed is detailed in Supplementary Results S2.7.

We analyzed the effect sizes over identified clusters using the Cohen’s d coefficient. Namely, for each site where a statistically significant effect of emotion or gaze was identified, we extracted the trial-by-trial activity within the time window of the cluster showing the largest sum of t-values. This activity was averaged across time, thereby obtaining one activity value per trial and per experimental condition, on each site. For two sample t-tests, Cohen’s d is computed as follows:

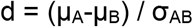

where μA-μB is the difference in the average activity for conditions A and B, and

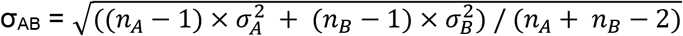

with σA, σB the standard deviations and nA, nB the number of trials for conditions A and B, respectively.

As the sign of d is dependent on the arbitrary polarity of the signal, we considered only absolute values of d. We interpreted the magnitude of Cohen’s d according to (Sawilowsky 2009). We performed two-sample t-tests across sites to compare the effect sizes for Gaze and Emotion in each ROI, and one-way ANOVAs followed by post-hoc t-tests to test for differences between the four ROIs on the effect size for Emotion and Gaze.

### Identification of healthy brain white matter tract endpoints relative to epilepsy patients’ intracerebral sites responsive to faces

Using the MNI coordinates of active bipolar sites identified in the epilepsy patients as a guidepost (see Supplementary Table 2 for active sites in individual patients), we attempted to identify overlap with the endpoints to various canonical posterior white matter tracts in occipitotemporal grey matter by interrogating the diffusion weighted imaging data of a large group of healthy non-epileptic subjects. The aim was to postulate potential routes of information flow in the brain that might account for the sequential and parallel information flow suggested by the latencies and amplitude differences of responses to faces between experimental conditions (Face 1 / Face 2) in our iEEG data.

#### Epilepsy patient intracerebral sites

MNI coordinates of active bipolar sites in each ROI, where responses to Face 1, Face 2, or both were observed (see previous section in Methods) were used for this analysis. Specifically, we considered the coordinates of the sites that responded to Face 1, including sites that responded either to Face 1 only, or to Face 1 and Face 2, and the coordinates of the sites that responded to Face 2 only. This decision followed the pattern of results that were obtained. Our rationale was to examine whether we could dissect the pathways for processing different attributes of the face stimulus, specifically those to viewing only facial motion/emotion (Face 2 only) versus other aspects of the face. We also included the coordinates from bipolar sites in the IPS (See Supplementary Results, S2.4) in this exploratory analysis.

#### Healthy brain data

##### Data sources

Diffusion, structural T1, and FreeSurfer segmentation (Fischl 2012) data were all obtained from the public release of the Human Connectome Project (Van Essen et al. 2013). For 1066 subjects (age range: 22 to 35 years), all three of these data modalities are hosted on the *brainlife.io* platform as a publicly accessible project (https://brainlife.io/project/5941a225f876b000210c11e5/detail). These HCP-provided data sources were subjected to the processing pipeline detailed below to obtain the tractography and segmentation data derivatives used in subsequent quantification and visualization.

##### Tractography

We performed anatomically constrained tracking (Smith et al. 2012), with tractography being generated for each subject using a *brainlife.io* implementation of mrtrix 3 (Fischl 2012; Tournier et al. 2019). This implementation is an openly available service (https://doi.org/10.25663/bl.app.101). The *brainlife.io* app implementation also incorporates ensemble tracking methods that perform a sweep across various turning, L_max_ and tractography parameters (Takemura et al. 2016) in order to generate a more accurate model of the human white matter. For this analysis we iterated across L_max_ parameters up to L_max_ 10, across maximum curvature parameters of 10, 20, 40 and 80 degrees, and performed both deterministic and probabilistic tracking. A minimum streamline length of 10 mm and a maximum streamline length of 200 were imposed, with a fractional anisotropy threshold greater than 0.2. A total of 25,000 streamlines were tracked for each combination of parameters, resulting in 600,000 streamlines being tracked per subject overall.

##### White matter tracts segmentation

Tractography segmentation was performed for each subject using an approach similar to White Matter Query Language (Wassermann et al. 2016). This method has been used previously to segment a number of underreported white matter tracts in posterior brain regions (i.e. pArc, TP-SPL, MdLF-Ang/SPL) (Bullock et al. 2019). Key to this approach is the use of cortical and subcortical anatomical landmarks to segment white matter tracts. Additionally, cleaning to remove streamline outliers (Yeatman et al. 2012) was performed so as to remove streamlines that were more than 4 standard deviations from the tract centroid or the average streamline length for the tract.

##### Cortical endpoint map generation

So as to ultimately plot the relation of endpoints to the location of active sites, it was necessary to generate an endpoint density mask for the end of each tract and each subject. This process began with reorienting the constituent streamlines of each tract such that they were oriented in the same direction. This reorientation was performed to ensure that the first and last node for each streamline corresponds to the appropriate endpoint collection (and thus all “first” nodes are computed with one another, and the same for “last” nodes). Subsequently, a count was performed for the number of first or last nodes in each 1 mm voxel (in native space). These outputs were smoothed with a 3 mm radius smoothing kernel. The resultant count information was stored as a nifti file.

##### Multi-subject map generation

In order to permit cross-subject comparison of tract endpoints, a warp to MNI space of the cortical endpoint maps was performed with ANTS (Avants et al. 2011) using a standard template (Fonov et al. 2011). Once endpoint maps had been warped to MNI space they were first thresholded at 0.01 endpoint density value, then binarized, and finally summed to obtain a count of the number of subjects exhibiting endpoints in each 1 mm voxel of the MNI volume.

#### Searching for overlap between tract endpoints and location of responsive sites

We investigated the overlap between the MNI co-ordinates of active bipolar sites from our patient group and tract endpoint masks derived from HCP subjects. Specifically, we sought to compute the overlap between each individual patient’s active site for each region of interest (FC, IOC, ITC, STC as well as the IPS) and endpoint masks from all segmented white matter tracts generated by a previously described, automated white matter segmentation method ((Bullock et al. 2019); https://brainlife.io/app/5cc73ef44ed9df00317f6288). This computation resulted in a pairwise proportion measure for each pairing of tract endpoint mask and electrode group, and thus formed an electrode group× tract endpoint mask data matrix. Importantly, only endpoint mask voxels containing endpoints from 100 or more subjects (i.e. 10%) were considered valid for the purposes of this computation. Thus, for each matrix entry (i.e. site-tract endpoint mask pairing), the numerical value could range from 0, indicating that no active site coordinate fell within the endpoint masks’ voxels, to 1, indicating that all of active site coordinates fell within the endpoint masks’ voxels. Tracts that exhibited no overlap with any site were excluded from the visualization (Fig. 9).

All ROIs containing active sites were found to exhibit overlap with at least one tract endpoint mask, and thus all were included in the visualization. Conversely, only a subset of the tract endpoint masks exhibited overlap with ROIs containing active sites and were thereby included in the visualization. We provide the complete code pipeline for performing this analysis and generating the associated figure in an open repository (https://github.com/DanNBullock/EcogAnalysisCode).

## RESULTS

### 1. ERP data: Response profile to face onset and social cue change

Here, we answer the first questions that we posed in the introduction: Where are the predominant sites that respond to face onset and changes in gaze and emotion across the face network? How does waveform morphology, amplitude and latency alter as a function of these facial attributes? For this, we compared responses to ‘Face 1’ and ‘Face 2’ stimuli. Face 1 represents the onset of a neutral direct gazing face. Face 2 corresponds to the dynamic change in socio-emotional facial expression. We focused our analyses on four regions of interest (ROI): the Inferior Occipital Cortex (IOC), the Fusiform Cortex (FC), and the Superior Temporal Cortex (STC), and the Inferior Temporal Cortex (ITC). Overall, across these 4 ROIs, we analyzed a total of 323 sites in the 11 patients (Fig. 2; Supplementary Table 2).

#### 1.1. Responsiveness to face onset (Face1) and changes in gaze and emotion (Face 2)

In each ROI, the proportion of responsive contacts for Face 1 and/or Face 2 was computed. This was larger in the right than the left hemisphere for all ROIs. However, as the number of sites was smaller in the left than in the right hemisphere, we cannot exclude that this may be due to less extended sampling of the left as compared to the right occipito-temporal regions. Responsiveness to each face type and the location of active bipolar sites is depicted in Figure 3 (see also Supplementary Table 2 and Supplementary Results S2.3 for more details).

**Figure 3.**
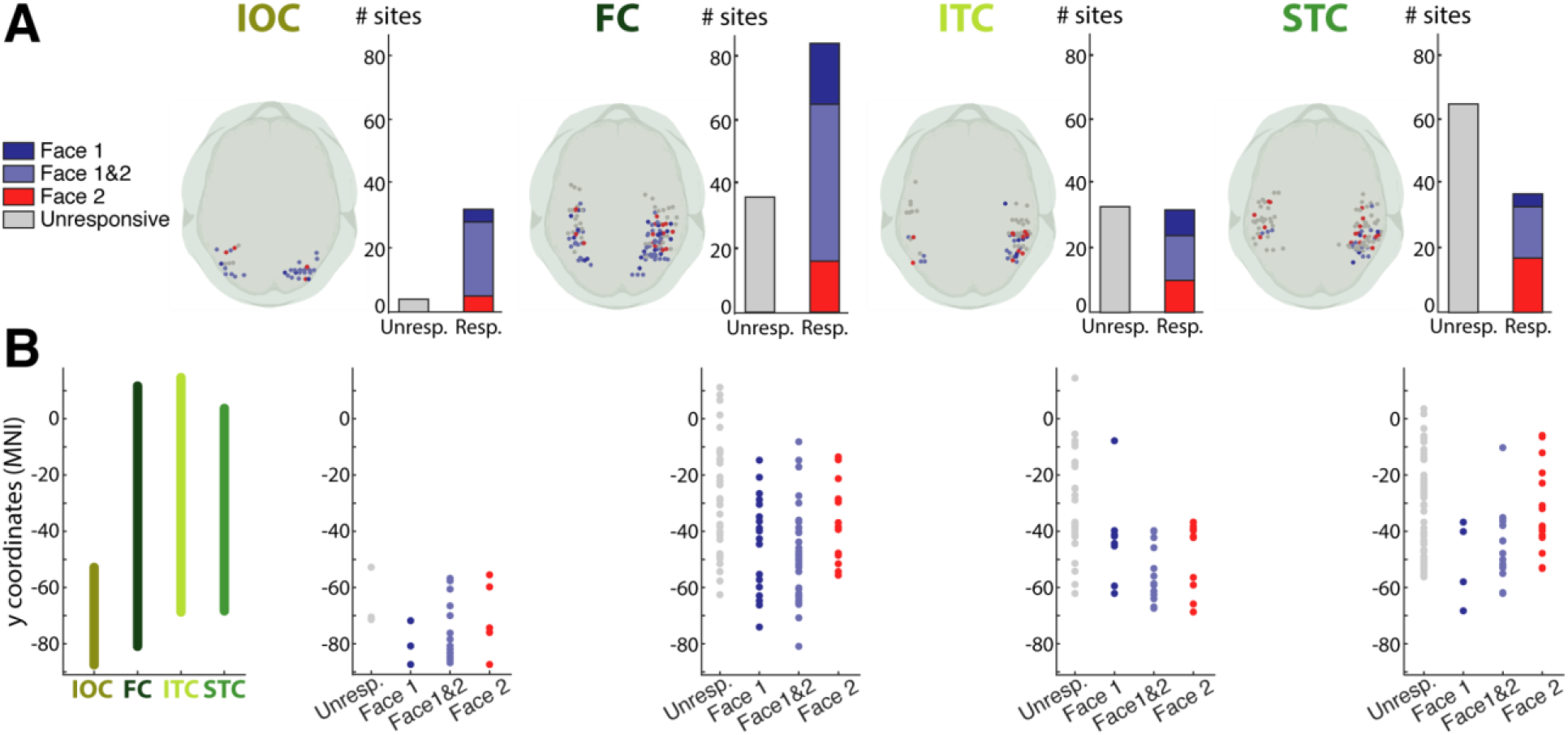
Responsiveness to face onset (Face 1) and change (Face 2) in the four regions of interest. **A. Number of responsive bipolar sites in each ROI.** The locations of the responsive sites to Face 1, Face 2, or both are represented on 3D axial brain view at the left of each ROI bar plot. For each ROI, the bar plots represent the total number of unresponsive sites (in grey), the number of sites responding only to Face 1 (in dark blue), only to Face 2 (in red), and to both Face 1 and Face 2 (in lavender). The responsiveness profile varied across ROIs. The IOC was the most responsive region and the STC showed a relative preference for Face 2 (See Supplementary Results S2.3 for details). **B. Distribution of unresponsive and responsive sites along the posterior-anterior axis.** The left plot indicates the posterior-anterior span across the y-coordinate of each of the 4 ROIs. The remaining four plots represent the location (y-coordinate) of the unresponsive and responsive sites in each ROI, pooling together the right and left hemisphere sites. Each color-coded dot represents a site, color-coded as in A. IOC, Inferior Occipital Cortex; FC, Fusiform Cortex; ITC, Inferior Temporal Cortex; STC, Superior Temporal Cortex.

**Figure 4.**
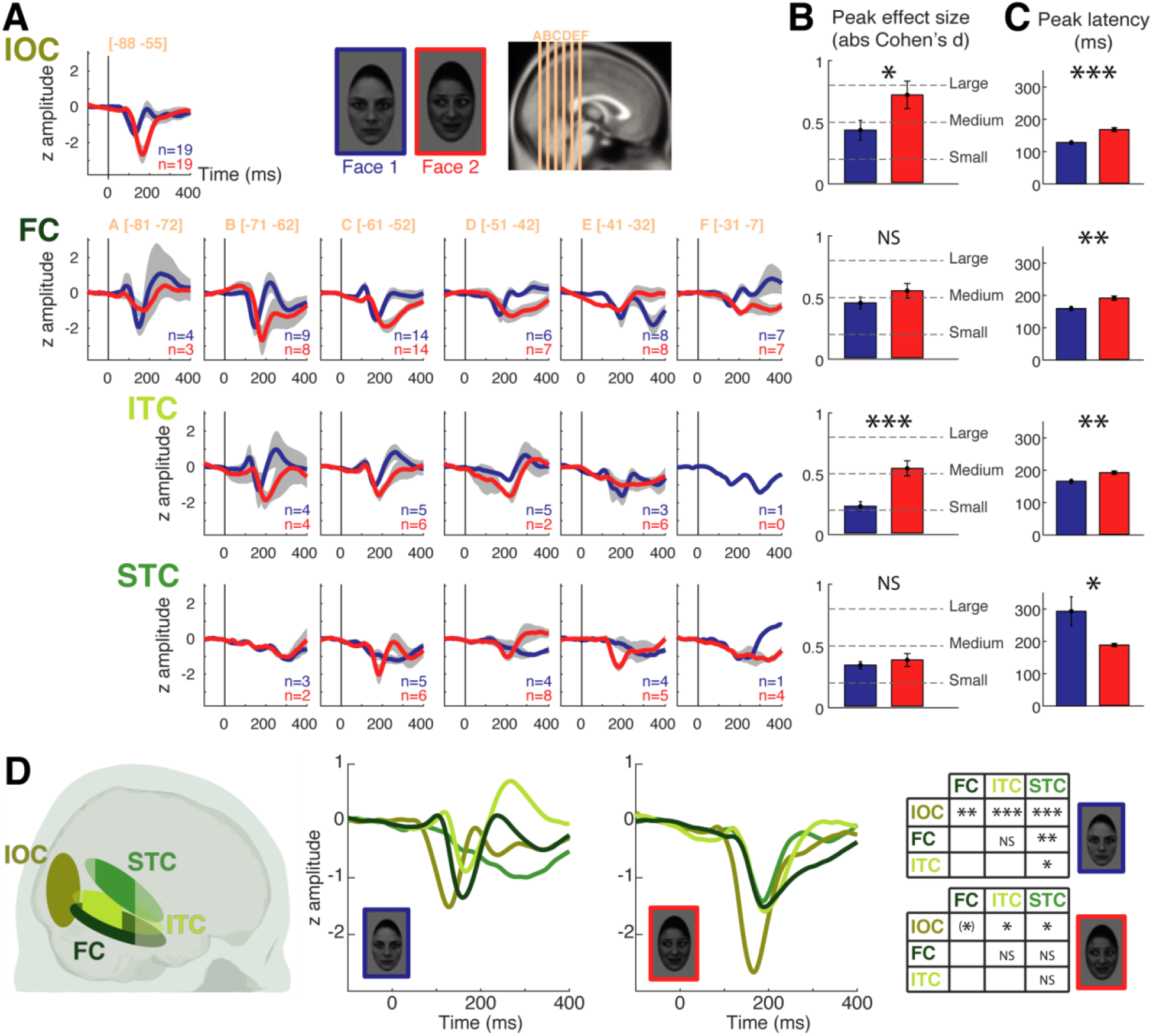
ERPs to Face 1 and Face 2 across the four ROIs in the right hemisphere. A. ERP morphology. ERPs in response to Face 1 (in blue) and Face 2 (in red) were averaged across patients, within each ROI, for each slice (A to F) along the posterior-to-anterior axis (for FC, ITC, STC). Bipolar sites for IOC were clustered in a narrow y-range and were therefore averaged altogether. The number of sites (n) averaged for each ERP is indicated on the bottom right-hand corner of each plot. The grey area around ERP time courses represents the standard error of the mean (no grey area for ERPs obtained from a single site). The amplitude of ERPs is expressed in z-scores and the scale is the same for all plots. See Supplementary Figures 2 to 5, for a complete illustration of all sites included in each slice. **B. ERP amplitude.** The absolute effect size (Cohen’s d) of the N200 to Face 1 (blue) and Face 2 (red) is shown for IOC (1st row), FC (2nd row), and ITC (3rd row); for STC (4th row), effect size was computed over the maximum of the slow ERP deflection to Face 1, while the peak of the N200 was considered for Face 2. We computed effect sizes over the slices showing the clearest ERPs (IOC: all sites; FC: slices A to E; ITC: slices B to D; STC: slices C to E). For each ROI, we statistically tested the difference in effect sizes between Face 1 and Face 2. The error bars represent the standard error of the mean. The dotted lines represent commonly accepted evaluations of the effect size measure. **C. Peak latency.**The latency of the N200 to Face 1 (blue) or Face 2 (red) is represented for all ROIs. As in B, it was computed from the slices showing the clearest ERPs. The latency computed for ERPs to Face 1 in the STC corresponds to the latency of the maximum of the slow ERP deflection. For each ROI, we statistically tested the difference in peak latency between Face 1 and Face 2, using a jack-knife procedure. Error bars represent the standard error of the mean derived from this procedure. **D. ERP comparison across ROIs.** A color-coded schematic representation of the location of the four ROIs is presented on the 3D sagittal view (left). The areas appearing in more intense color correspond to the slices where ERPs were analyzed (see B). The middle two panels depict ERP waveforms to Face 1 (left) and to Face 2 (right) that were obtained by averaging ERPs in each ROI (taking into account the slices with the clearest ERPs as in B and C). In the rightmost display panel, two tables (for Face 1 and Face 2 respectively) show the results of the statistical comparisons of peak latencies across each pair of ROIs. IOC, Inferior Occipital Cortex; FC, Fusiform Cortex; ITC, Inferior Temporal Cortex; STC, Superior Temporal Cortex. NS: non-significant; (*): p<0.08; *: p<0.05; **: p<0.01; ***: p<0.005.

The responsive sites occurred over a large posterior-to-anterior portion for the FC (y = − 81 to −8) (Fig. 3B). In ITC, most responsive sites were found posteriorly (y = −69 to −37, with only one responsive site located anteriorly: y = −8); anterior sites did not appear to be responsive to either Face 1 or Face 2. The STC showed a gradient in responsiveness type, where sites responding to Face 1 were mostly found in posterior portions of STC (y = −68 to −35, with only one site outside this range, y = −10, which responded to both Face 1 and Face 2) and sites responding selectively to Face 2 were present in more anterior portions of the STC (y = −53 to −6).

#### 1.2. Waveform morphology, amplitude, and latency

We characterized specific ERP attributes, i.e., morphology, amplitude, and latency, focusing the analysis on the right hemisphere where we had more extensive sampling. For the 3 ROIs with the largest y-range (FC, ITC, and STC), ERP morphology changed along the posterior-to-anterior axis, with the ‘typical’ morphology occurring at intermediate points on this axis and smaller and more variable responses occurring at the posterior and/or anterior borders (Fig. 4A). This inhomogeneity of responses across the y-axis led us to further divide the FC, ITC, and STC regions into successive coronal slices of 10 mm along the posterior-to-anterior axis for the purposes of ERP morphology visualization and characterization.

##### ERP morphology

ERPs to both Face 1 and Face 2 in IOC, FC, and ITC were typically characterized by a sharp, large amplitude negative deflection peaking between 150 and 200 ms (Fig. 4A, see also Supplementary Figures 2 to 5 for an illustration of individual data in each ROI). This deflection corresponded to the N200 that has been previously described (together with other components; (Puce et al. 1999)), although it appeared to peak earlier in the IOC to Face 1. N200 was followed by a positive deflection (P250) of smaller amplitude, which was most prominent in response to Face 1. Additionally, in FC and ITC, there was a small positive and early deflection, corresponding to a P100. In the STC, ERPs to Face 2 also showed a stereotypical and sharp N200. This was in stark contrast to ERPs to Face 1, which were in the shape of a slower wave reaching its maximum at around 300 ms (seen most clearly in Fig. 4D).

##### ERP amplitude and latency

We then determined peak amplitude and latency of the main negative deflection in response to Face 1 and Face 2, in each ROI. In the ITC, FC, and STC, we concentrated on the slices where this response could be clearly identified (ITC: slices B to D; FC: slices A to E; STC: slices C to E; see Methods and Fig. 4A).

To compare the magnitude and reliability of ERPs between conditions, we computed peak amplitude in the form of *effect sizes* for the maximal early negative deflection in each ROI (that is, N200, except for STC responses to Face 1 where the maximum of the slow wave was selected) (Fig. 4B). N200 to Face 2 was significantly larger than that to Face 1 in IOC (two-sample t-test, t(36)=−2.09, p=0.044) and in ITC (t(24)=−4.36, p=0.0002). There was no significant amplitude difference between the N200 to Face 1 and 2 in FC (t(79)=−1.31, p=0.19). There was also no amplitude difference between the slow wave to Face 1 and the N200 to Face 2 in the STC (t(30)=− 0.61, p=0.55). These results were confirmed by an additional analysis using linear mixed effects models (see Supplementary Results S2.6), allowing to take into account inter-patient and inter-contact variability.

With respect to latency, the jackknife procedure indicated that the N200 peaked earlier to Face 1 than Face 2 for IOC (t(36)=4.32, p=0.0001, 95% confidence interval [CI] of the latency difference: [20 55 ms]), ITC (t(24)=3.02, p=0.0059, CI=[8 42 ms]), and FC (t(79)=3.03, p=0.0033, CI=[10 50 ms]) (Fig. 4C). Notably, the opposite pattern was observed for STC, where N200 to Face 2 peaked well *before* the maximum amplitude occurred to Face 1 (t(30)=−2.54, p=0.016, CI=[−207 −23 ms]).

When examined across ROIs, N200 showed a significant difference across ROIs for Face 1 (ANOVA on jack-knife latencies: p=0.0088). N200 to Face 1 peaked *first* in the IOC (Fig. 4D; jack-knife procedure with two-sample t-tests between each pair of ROIs: IOC vs. ITC: p=0.0011, IOC vs. FC: p=0.0060, IOC vs. STC: p=0.0008; Bonferroni corrected for multiple comparisons across the six 2-by-2 tests), followed by ITC and FC (ITC vs. FC: p=1), and lastly by the STC (STC vs. ITC: p=0.013; STC vs. FC: p=0.0052). For Face 2, the latency difference among ROIs was less marked, with a non-significant overall difference across ROIs (ANOVA on jack-knife latencies: p=0.26). There was no significant difference in N200 latency among ITC, STC, and FC (all p=1); the response to Face 2 was however marginally faster in IOC relative to ITC, STC, and FC (IOC vs. ITC: p=0.027, IOC vs. STC: p=0.046, IOC vs. FC: p=0.054; all p-values Bonferroni corrected). Using linear mixed effects model analyses, we confirmed these effects and directly probed the sequential order of peak latencies suggested here; this allowed confirming that ERPs in IOC were the earliest ones for both Face 1 and Face 2 (see Supplementary Results S2.6).

#### 1.3. Interim summary: Response profile to face onset and social cue change

In sum, we observed consistent responses to face onset (Face 1) and to face changes (Face 2), distributed along the posterior-to-anterior axis of the four ROIs. From our data, the IOC showed the earliest N200 peak latencies, for both Face 1 and Face 2. For Face 1, a sequence of activation from IOC, to ITC/FC and subsequently to STC emerged, as indicated by progressively increasing N200 peak latencies. Notably, in the STC, the activity was quite late, broad and slurred relative to the other three regions. In contrast, in response to Face 2, prominent N200 activity in IOC was followed by the parallel activation of ventral and dorsal pathways (FC, ITC and STC) as indicated by comparable N200 peak latencies. In this case, evoked activity in the STC to Face 2 was markedly different to that of Face 1, with a clear, sharp N200, similar to that from the other regions.

### 2. Sensitivity to gaze and emotion changes

We then turned to our next question: What parts of the face network are sensitive to changes in gaze direction vs. emotional expression?

#### 2.1. Comparative sensitivity to social cues across ROIs

To investigate the sensitivity to social cues, that is, the effects of emotion (happy / fearful) and gaze (averted / direct), we tested the ERPs at the sites that were significantly responsive to Face 2, in the 4 ROIs. As a preliminary step, we evaluated the proportion of sites that showed any statistically significant effect of gaze, emotion, or emotion-by-gaze interaction within each ROI (see Supplementary Results S2.7).

To identify where we had the larger signal-to-noise ratios in response to gaze and emotion, we examined *effect size* at each site and each ROI (Fig. 5). No statistically significant difference between effect sizes for gaze and emotion was found in either IOC, FC, or ITC (two-sample t-tests: t(17)=0.75, p=0.46 for IOC; t(28)=0.63, p=0.53 for FC; t(8)=−2.01, p=0.079 for ITC), where the effect sizes were small to medium for both social cues. In contrast, there was a marked difference in effect size for *gaze* relative to emotion in STC (t(13)=−3.04, p=0.0096). The mean effect size was small to medium for emotion whereas it was medium to large for gaze. As a consequence, the size of gaze effects differed significantly between the four ROIs (F(3,24)=6.38, p=0.0025), with greater gaze effect size in STC than in IOC (t(14)=−2.95, p=0.010), FC (t(16)=− 3.11, p=0.0067), and to a lesser extend ITC (t(10)=−2.02, p=0.071). In contrast, the size of emotion effects did not differ across the ROIs (F(3, 42)=2.12, p=0.11). These results were further confirmed by linear mixed effects model analyses, taking into account inter-patient variability (see Supplementary Results S2.8).

**Figure 5.**
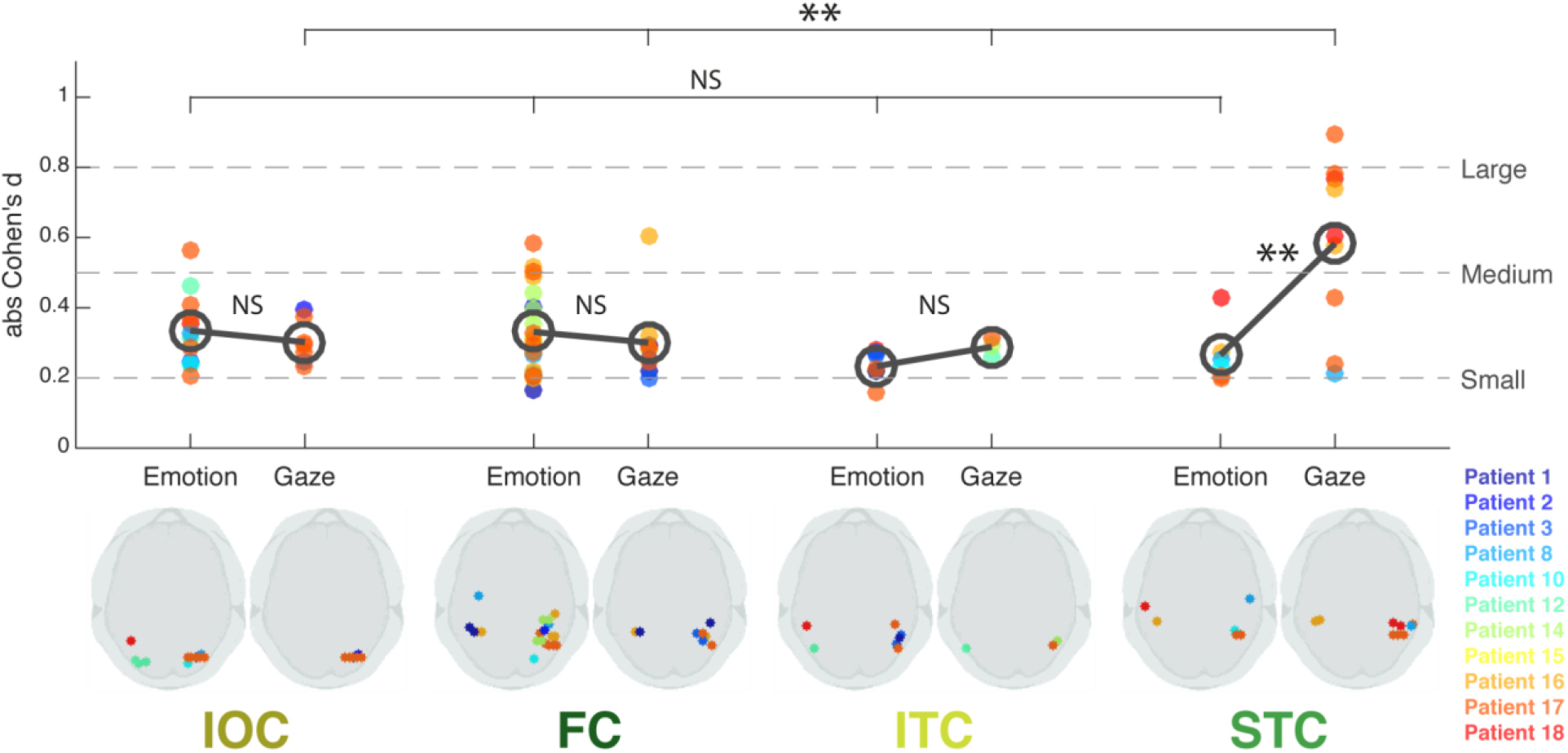
Effect Sizes for Emotion and Gaze for each of the four ROIs. For each ROI (IOC, FC, ITC, and STC), each dot corresponds to one bipolar site, significantly responding to Emotion (on the left) or Gaze (on the right), with the corresponding effect size (absolute Cohen’s d) represented on the y-axis. The dark grey open circle represents the mean effect size across sites, for Emotion and Gaze respectively, within each ROI. Effect sizes were compared between Emotion and Gaze in each ROI (grey bars between the open circles) and across ROIs for each Emotion and Gaze effect (top grey bars). The dotted lines represent commonly acceptable evaluations of the effect size values. The responsive site location can be viewed on the 3D axial brain silhouette views, for each effect in each ROI (bottom row), and the corresponding patient number is identifiable with the dot color (see color-patient correspondence on the bottom right). IOC, Inferior Occipital Cortex; FC, Fusiform Cortex; ITC, Inferior Temporal Cortex; STC, Superior Temporal Cortex. NS: non-significant; *: p<0.05; **: p<0.01.

In the next sections, we examine in more detail the responses to each social cue (gaze and emotion changes) in the 4 ROIs, in each patient.

#### 2.2. Responses to gaze in individual patients

Gaze effects were reliably found within the STC in 4 patients (Fig. 6). They were observed in a restricted portion of the STC, between y=−53 and y=−35, in both left and right hemispheres. Notably, the majority of the sites that showed a significant effect of Gaze were located in the Superior Temporal Sulcus (STS, 6 sites in right STS and 2 in left STS, 1 in right mid-temporal gyrus). In Figure 7, we display the results of patient 17, who had an extensive electrode sampling and showed significant gaze effects in all four ROIs. Specifically, in this patient we observed a double polarity reversal over three consecutive sites along an electrode in the right STS (Electrode 5 sites 1-3). These sites were located in the right posterior STS (y = −53), and sampled medial (x = 43, site 1) to more lateral (x = 54, site 3) parts of the sulcus. This double polarity inversion is a clear sign of high anatomical specificity, that is, of a local source, for the Gaze effect. These Gaze effects were observed between 130 and 400 ms, with larger N200 amplitudes for averted gaze, followed by a larger P250 in both the most medial and the most superficial sites. In contrast, the ERPs to direct gaze were small to negligible. This resulted in medium (|d|=0.43) to large (|d| > 0.77) gaze effect sizes across sites for this patient.

**Figure 6.**
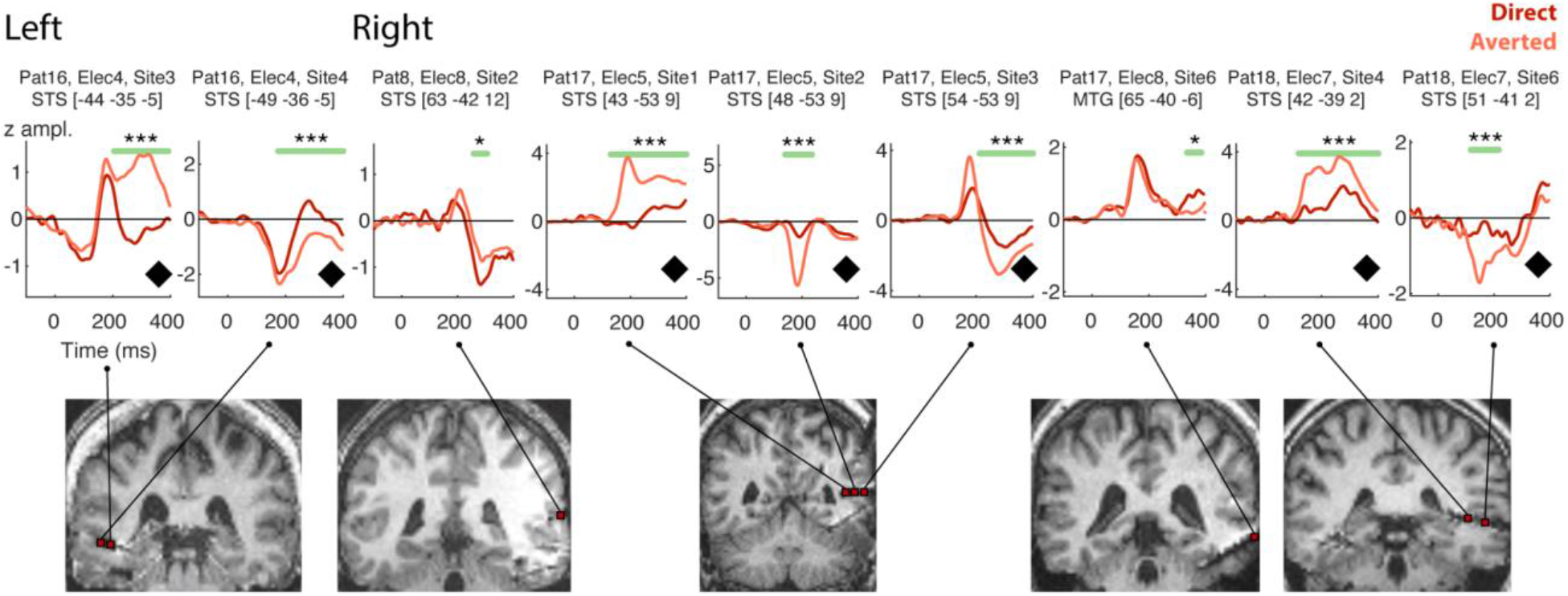
ERPs from sites in the Superior Temporal Cortex region (STC) responding to Gaze. The patient, electrode and site number, as well as the anatomical localization and the MNI coordinates are indicated for each ERP. Horizontal green bars indicate the time window where the conditions significantly differ. The coronal MRI figures show the localization of each site, on the patient’s normalized brain, at the corresponding y-coordinate. Black diamonds near x-axis indicate polarity inversions in adjacent sites. MTG: Mid-Temporal Gyrus, STS: Superior Temporal Sulcus. *: p<0.05, ***: p<0.005, Monte-Carlo p-values.

**Figure 7.**
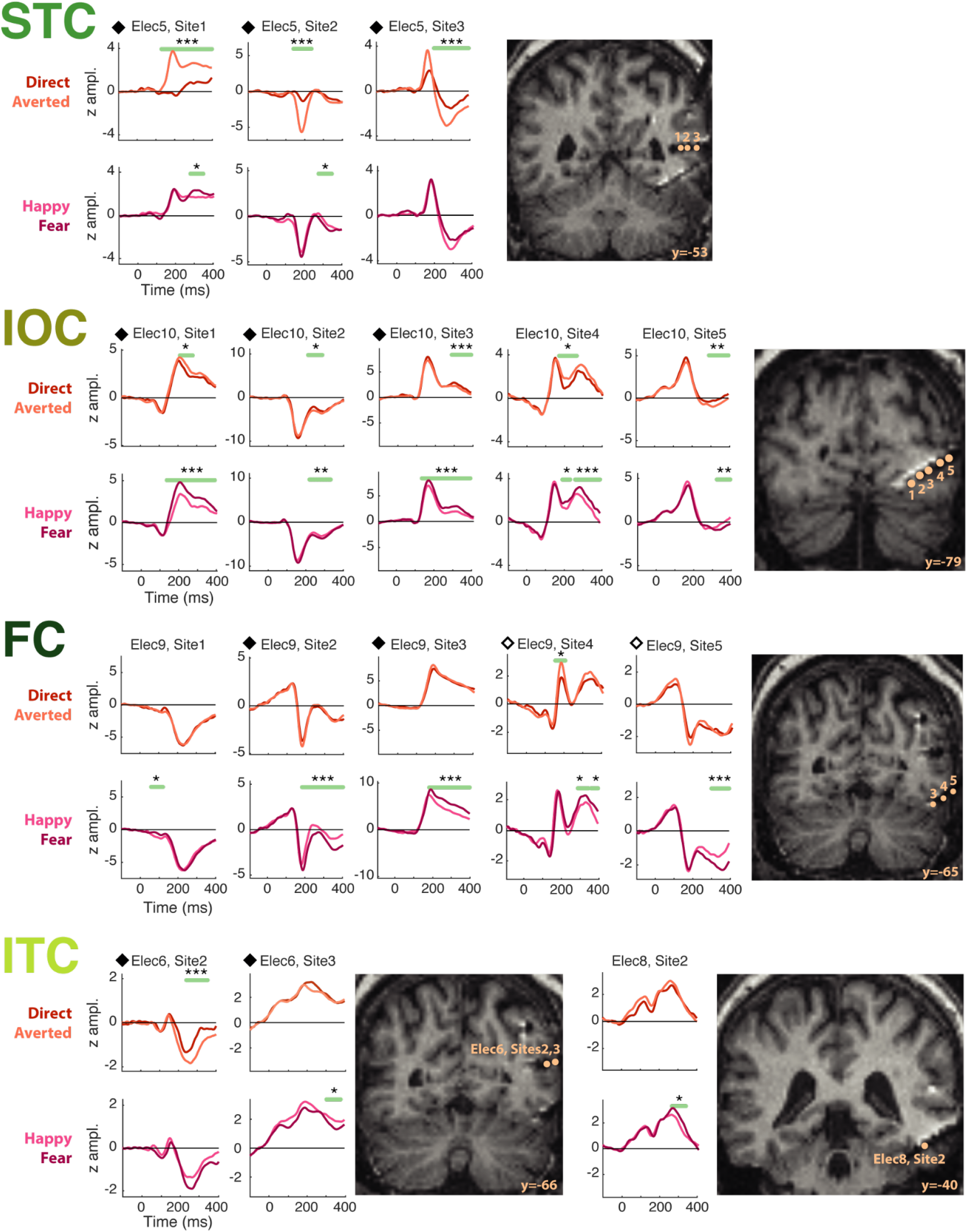
Effects of gaze and emotion within the four ROIs in Patient 17. This patient was selected due to extensive occipitotemporal sampling and because he displayed significant effects across all four ROIs; we selected representative electrode shafts in each ROI for this patient in order to illustrate both the effects of gaze and the effects of emotion (see Supplementary Fig. 8 to 11 for all effects, over all patients). The STC showed the most marked differences between averted and direct gaze conditions (top line of plots), with larger ERPs for averted gaze. The fearful vs happy emotion conditions produced subtle but quite long-lasting ERP changes that could occur across the entire ERP time course, at least in the FC and IOC of this patient. The sites are projected on coronal views of the patient’s post-implantation structural MRI. Note that sites 1 and 2 of Elec 9 in FC are not visualized because they were located in coronal slices different from sites 3-5. Black (solid and open) diamonds on top of the y-axis of the plots indicate polarity inversions between adjacent sites. ERP polarities were not rectified in these plots, in order to visualize polarity inversions. FC, Fusiform Cortex; IOC, Inferior Occipital Cortex; ITC, Inferior Temporal Cortex; STC, Superior Temporal Cortex. *: p<0.05, **: p<0.01, ***: p<0.005, corrected-over-time Monte-Carlo p-values.

Patients 16 and 18 also showed marked gaze effects, with polarity reversals within the STS (Fig. 6), more anteriorly than in Patient 17 (Patient 16: Electrode 4 sites 3-4; Patient 18: Electrode 7 sites 4 and 6), but in different hemispheres (MNI coordinates of mid-point between sites with polarity inversions, for Patient 16: −47 −36 −5; for Patient 18: 47 −40 2). Gaze effects occurred with large effect sizes (Patient 16: |d| > 0.57; Patient 18: |d| > 0.60) in similar time windows within 200 to 400 ms in both patients. A fourth patient (Patient 8) showed a more nuanced effect and later effect of gaze (|d| = 0.2, statistically significant effect between 260 ms and 300 ms).

In contrast, Gaze effects were smaller in the other ROIs (Fig. 5) and corresponded to more subtle modulations of ERP amplitude (Fig. 7, Supplementary Fig. 8-11). This can be notably seen in sites presenting a polarity inversion, hence revealing a high spatial specificity, e.g., within the IOC for Patient 17 (Fig. 7, Electrode 10 sites 1-3) and within the FC for patient 16 (Supplementary Fig. 8, Electrode 7 sites 1-2). The Gaze effects on these sites were much smaller in size (Patient 17: |d| < 0.30 across the IOC sites with polarity inversion; Patient 16: |d| < 0.33 across the FC sites with polarity inversion) than the large, also local, STC effects observed in those same patients.

We made sure that our gaze effects were not confounded by emotion, which could potentially be the case due to an imbalance in the number of trials (see Supplementary results S2.10). Furthermore, to evaluate potential effects of the direction of gaze change, we tested all the sites showing a significant effect of gaze change for differences between leftward and rightward gaze change conditions (see Supplementary results S2.11).

#### 2.3. Responses to emotion in individual patients

The effects of emotion were more subtle than those observed for gaze. In general, they consisted of amplitude modulations of the ERP, as shown in Figure 7 for Patient 17 (see also Supplementary Fig. 8-11). Here, we looked at how emotion and gaze were processed in the sites that were sensitive to both social cues.

In the STC, there were three such sites (Supplementary Fig. 10; Patient 16: Electrode 4 site 4; Patient 17: Electrode 5 sites 1-2). In agreement with the finding of a specific gaze effect in this ROI, the emotion effects on these sites were smaller and of shorter duration than the gaze effects and occurred within the same time window or later on. In addition, one STC site of Patient 18 (Electrode 7 site 4) showed a late, short-lived interaction between gaze and emotion, embedded within a long-lasting and large gaze effect. Altogether, these data confirm that STC prioritizes gaze rather than emotion processing.

In the FC (Supplementary Fig. 8), a total of 21 sites responded to emotion, and among these, three sites responded to both emotion and gaze (Patient 16: Electrode 7 sites 1-2; Patient 17: Electrode 9 site 4), in different and non-overlapping time windows. Patient 17 also had a FC site (Electrode 9 site 2) responding to emotion over a long time window (~200 to 400 ms), together with a short-lived interaction between emotion and gaze at ~300 ms.

In the IOC (Supplementary Fig. 9), six sites showed both emotion and gaze effects (Patient 3: Electrode 7 site 2; Patient 17: Electrode 10 sites 1-5), in overlapping time windows. These effects were short-lasting and small in Patient 3. In contrast, Patient 17 showed remarkably larger and longer effects for emotion than gaze.

In the ITC (Supplementary Fig. 11), no site showed both emotion and gaze effect, however, one site (Patient 17: Electrode 6 site 2) showed some interaction effects, partly embedded within the time window of a gaze effect.

#### 2.4 Interim summary: Sensitivity to gaze and emotion changes

Responses to gaze and emotion changes were present in all four ROIs. In the STC, and specifically the STS, we observed the largest effect sizes to the gaze change, with a striking increase of ERP amplitude for averted relative to direct gaze. This result was evident across different bipolar sites from different patients. Accompanying polarity inversions suggested a local generator for this activity (Fig. 6, 7). The effects of emotion were generally more subtle.

### 3. Healthy brain white matter tract endpoints relative to epilepsy patient iEEG sites active to faces

We next turned to the white matter tract endpoint analysis to answer our last question: What are the likely routes of information flow across structures in the face network?

We identified the cortical projection zone of a series of posterior white matter tracts [in healthy brains] and estimated potential overlap with each measurement in the iEEG data. The relevant white matter tracts connecting the dorsal, ventral and lateral posterior human cortex included the vertical occipital fasciculus (VOF; Yeatman et al. 2014; Takemura et al. 2017), the arcuate fasciculus (Arc) and posterior arcuate (pArc) (Catani and Mesulam 2008; Weiner et al. 2017), the temporo-parietal connection (TP-SPL), the middle longitudinal fasciculus (MdLF; Bullock et al. 2019). Other tracts of interest (spanning the anteroposterior brain axis) were the inferior longitudinal fasciculus (ILF), superior longitudinal fasciculus (SLF).

#### 3.1 Healthy brain white matter tract endpoints

We first calculated and visualized the endpoints of the abovementioned white matter tracts in 1066 HCP healthy subjects in MNI coordinate space. The endpoints of some of these structures are displayed on inflated cortical surfaces as a function of overlap in the HCP subjects (Fig. 8). Some endpoints had an extensive and more variable distribution across subjects, e.g. the dorsal aspect of the Arc, the superior parietal lobule (SPL) endpoints of the MdLF, i.e. MdLF-SPL, whereas others had a much more homogeneous distribution across individuals, despite still having a substantial cortical footprint— e.g., the ventral aspect of the VOF, anterior temporo-polar aspect of ILF, and posterior aspects of the subcomponents 1 and 2 of the superior longitudinal fasciculus (SLF) (Thiebaut de Schotten et al. 2011).

**Figure 8.**
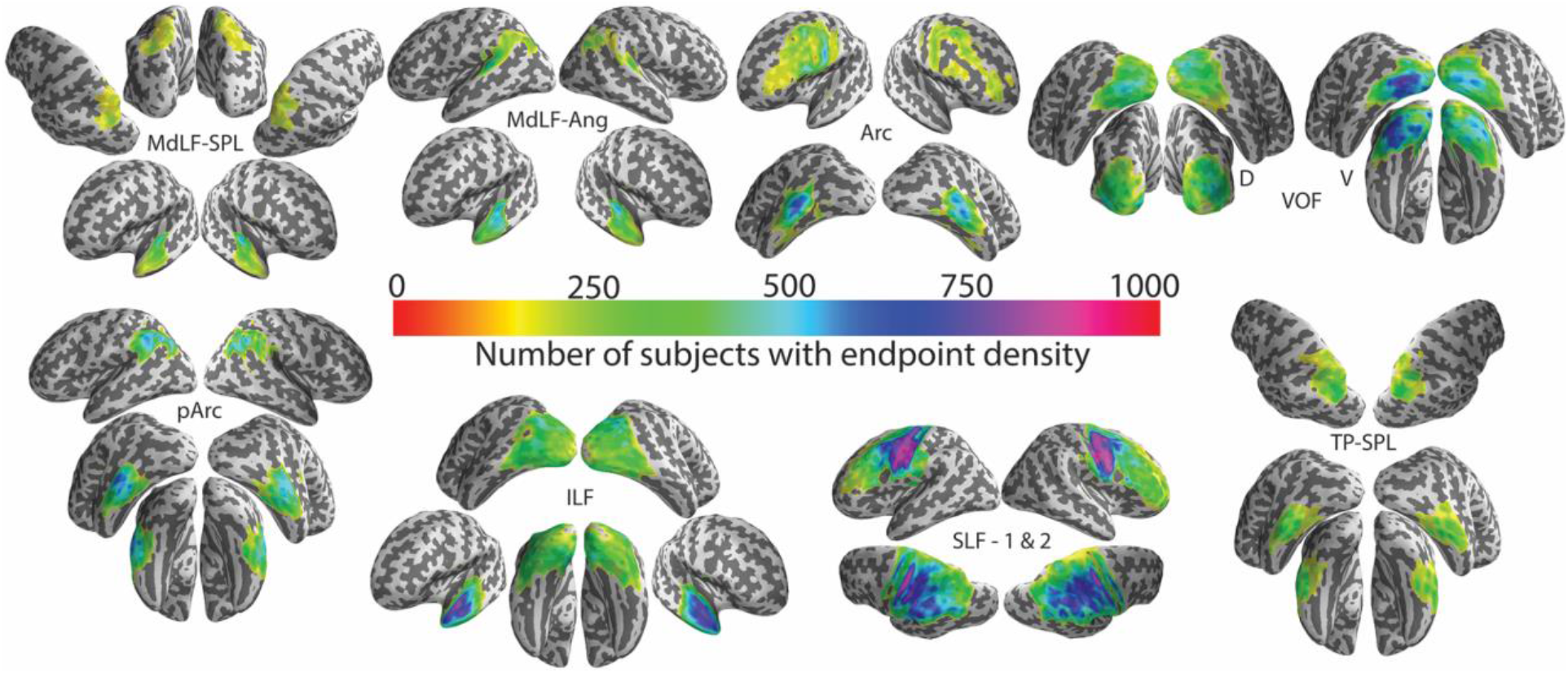
Endpoints for canonical white matter tracts that course partly, or wholly, through occipitotemporal regions. A series of inflated cortical surfaces display grey matter endpoints for the middle longitudinal fasciculus (MdLF) of the superior parietal lobule (SPL) and the angular gyrus (Ang), arcuate fasciculus (Arc), vertical occipital fasciculus (VOF, D=dorsal, V=ventral), posterior Arc (pArc), inferior longitudinal fasciculus (ILF), superior longitudinal fasciculus (SLF) subcomponents 1 and 2, the temporo-parietal connection of the superior parietal lobule (TP-SPL). Color scales display the number of subjects showing voxels at these endpoints in a density histogram that ranges from 1 to 1000 (red through to pink). Data have been thresholded at 150.

#### 3.2 Overlap of white matter tract endpoints and active iEEG sites

The overlap analysis between the computed white matter tract endpoints and the coordinates of active bipolar sites appears in Figure 9. Based on the calculated proportions for sites responding to Face 1 and to Face 2, the most overlap occurred for the more posteriorly located ROI of the IOC in the posterior endpoints of ILF. IOC active sites also overlapped to lesser extent with the VOF. Interestingly, IPS sites also intersected with the posterior endpoints of ILF, possibly abutting the posterior occipito-parietal arch of this tract. Furthermore, IPS active sites showed some overlap with the posterior end of subcomponents 1 and 2 of the SLF and of MdLF-SPL. For the more anteriorly located ITC sites, the inferior end of Arc, pArc and TP-SPL white matter tracts were prominent overlap points. There was also a scarce overlap with anterior ILF, likely associated with the most anterior active ITC site. An overlap with inferior Arc, pArc and TP-SPL endpoints was also observed for the FC sites, with additional overlap with posterior ILF. Finally, for the STC, active sites overlapped with inferior endpoints of pArc and Arc and posterior endpoints of SLF, with additional scarce overlap with inferior TP-SPL, superior pArc and posterior MdLF-Ang.

**Figure 9.**
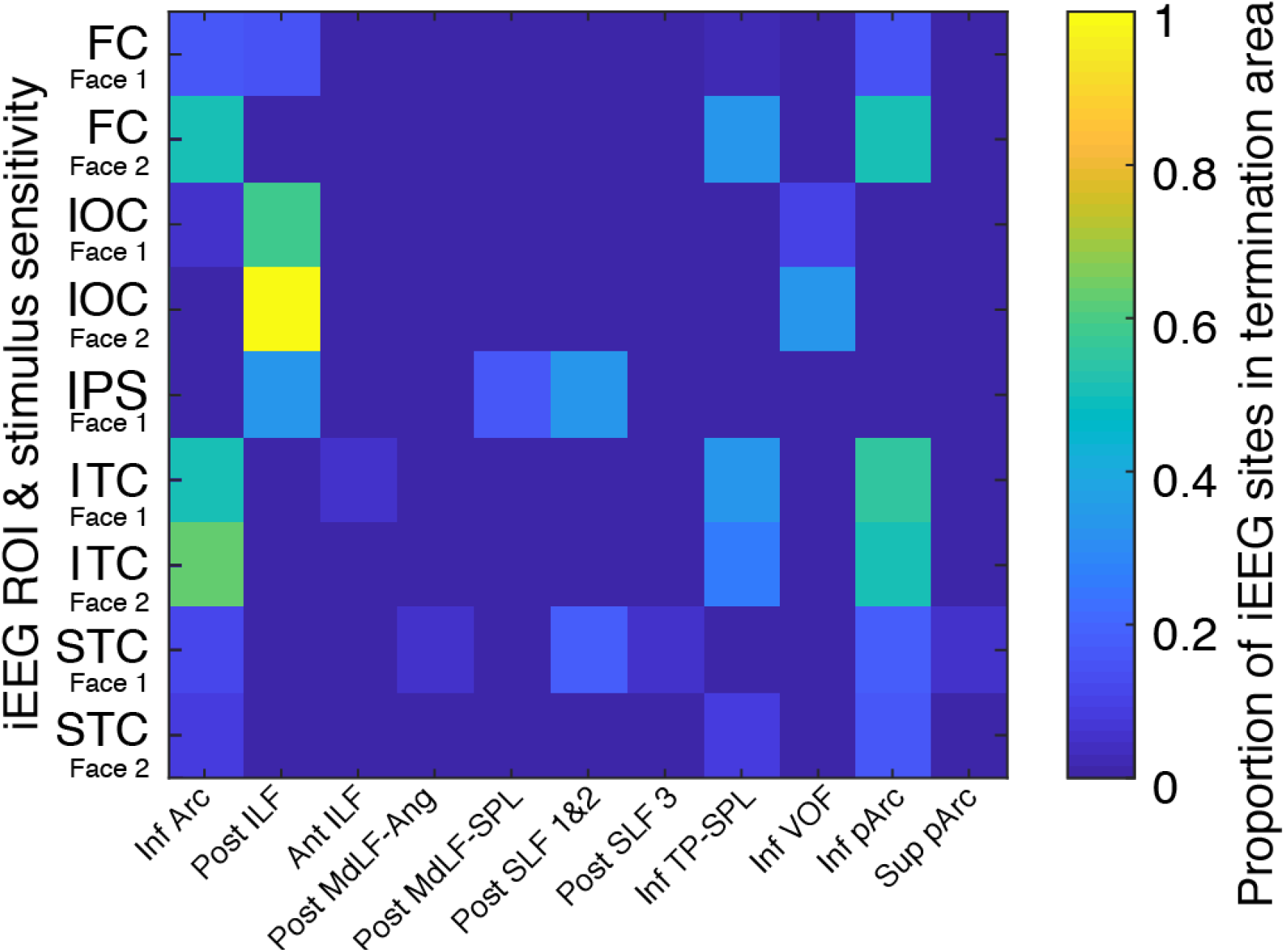
Overlap analysis between posterior white matter tract endpoints and active bipolar iEEG sites. The rows in the matrix are the ROIs (IOC, ITC, FC, STC, and IPS), with one row for the sites that responded to Face 1 (including the sites that responded to Face 1 only and the sites that responded to both Face 1 and Face 2, see Methods; labelled as Face 1) and one row for the sites that responded to Face 2 only (labelled as Face 2), for every ROI except IPS where no site was found to respond to Face 2 only. The columns are the white matter tract endpoints. The color scale indicates the proportion of the active sites that lay in a given tract endpoint zone. LEGEND: Inf, Sup, Post, Ant indicate the inferior and superior parts or the posterior and anterior parts (respectively) of the indicated tracts; arcuate fasciculus (Arc), inferior longitudinal fasciculus (ILF), angular gyrus and superior parietal lobule sections of the middle longitudinal fasciculus (MdLF-Ang and MdLF-SPL, respectively), superior longitudinal fasciculus (SLF) [where 1, 2 and 3 denote the 3 subcomponents of the SLF, see (Thiebaut de Schotten et al. 2011)], temporo-parietal connection of the superior parietal lobule (TP-SPL), vertical occipital fasciculus (VOF), and posterior arcuate (pArc).

## DISCUSSION

Here we investigated face processing using iEEG recordings from 323 bipolar sites in the occipito-temporal cortex of 11 patients. We first examined responsiveness to the onset of a neutral face (i.e. Face 1) and a subsequent change of facial social cues (i.e. Face 2) in four anatomically defined ROIs—IOC, FC, ITC, and STC. We then characterized the *sequence of activation* in these ROIs using N200 latency. We further analyzed the effects of social cues (gaze and emotion changes) at the group and individual levels. Finally, we examined the white matter tracts that may underlie information flow across active sites. Several main results emerged from these analyses. First, the IOC consistently showed the earliest latencies for both face stimulus types, consistent with the claim that it could be an entry point for information flowing into the face-processing network. Second, STC responses to faces showed distinctive features, including clear modification by stimulus type. In particular, gaze/expression changes elicited significantly earlier ERPs relative to static face onset, while the *opposite was true* in IOC, ITC, and FC. The effect of gaze was also distinctive—showing greater effect size—in STC, in comparison to the effect of emotion and in comparison, to the effect of gaze in the other ROIs. We also observed that the ITC, a region that is not usually described as part of the face network, showed a vigorous response to all facial stimuli. We will discuss these results before turning to white matter tracts that may underlie the temporal unfolding of face and facial social cue processing.

### The IOC is a potential entry point into the core face processing network

Robust responses to the face onsets and changes were observed in the latency range of the N200 in the four ROIs. N200 latency was the earliest in the IOC ROI, as compared to the other ROIs—ITC, FC, and STC. This suggests that the IOC is likely to be the entry point into the network, consistent with some previous studies that were based on fMRI activation in healthy subjects, as well as neuropsychological lesion and non-invasive stimulation studies (Haxby et al. 2000; Rossion et al. 2003; Fairhall and Ishai 2007; Pitcher et al. 2007). The idea that IOC (in terms of the inferior occipital gyrus) provides input to the face processing network has been previously advanced (Haxby et al. 2000; Pitcher et al. 2007). Fairhall and Ishai (2007) provided some supporting evidence based on fMRI data for the processing of static famous faces (Fairhall and Ishai 2007), but other studies emphasized IOC as part of the ventral pathway of face processing, hence mainly in association with the FC (Gobbini and Haxby 2007; Pitcher et al. 2014; Pitcher, Pilkington, et al. 2020), putting into question the hierarchical nature of the core face network (Rossion, 2008; Atkinson and Adolphs, 2011). In the model for famous face recognition of Fairhall and Ishai, the IOC was thought to send its output to both the FG and the STS (Fairhall and Ishai 2007). This is consistent with our results where the shortest latencies were obtained in the IOC ROI for both face onsets and social cue changes and IOC showed greater response to social cue changes than face onsets, leading us to conclude that the IOC might be a likely entry point for facial information into both the ventral and dorsal face processing pathways. That said, two points need to be made. First, we did not have intracranial electrodes in earlier visual regions e.g. V1, V2 etc. so we cannot talk about the complete route that the visual information related to faces might take in the visual system. While IOC appears as an entry point of the core face network in our study, other studies have shown that it is not the sole entry point to the system, particularly in lesioned brain cases (Weiner et al., 2016). Second, the fact that IOC appears as an entry point of the core face network in our study does not posit it as a low-level processing region. It has been shown that IOC, including face-selective and not-face-selective regions, perform high-level face processing (Dricot et al., 2008; Avidan and Behrmann, 2009). In addition, it may be noted that in an fMRI study using face and cars embedded in noise, Jiang et al. (2011) concluded that the fMRI activity in OFA [likely to be in our IOC ROI] *lagged* that in the FFA. However, fMRI is likely to conflate neurophysiological activities that can be observed at early and late latencies in the very same cortical sites (e.g. Puce et al., 1999), making it difficult—if not impossible—to make parallels about timing from fMRI and electrophysiological studies.

Individual data analysis further indicated that IOC was sensitive to both emotion and gaze in overlapping time periods. In a combined transcranial magnetic stimulation (TMS)-fMRI study, TMS was given to either to the right occipital face area (OFA) or posterior STS (Pitcher et al. 2014). Interestingly, stimulation of the OFA reduced fusiform face area (FFA) activation to both static and dynamic faces and STS activation to static, but not dynamic faces. The authors suggested that the processing of dynamic information from faces may bypass the OFA, involving an adjacent region of movement processing (MT/V5). They have further recently proposed that the information from MT/V5 may be fed directly to the STS—forming a third visual pathway (Pitcher and Ungerleider, 2021). This is not incompatible with our results because our anatomically defined IOC ROI is likely to have encompassed both OFA and MT/V5 functional regions (Dumoulin et al., 2000). Indeed, we have previously observed face-related activity in MT/V5 in both fMRI (Wheaton et al., 2004) and MEG (Watanabe et al., 2001, 2006) studies. Data from the current study would be consistent with the finding that several IOC sites showed responses to both emotion and gaze in overlapping time periods, with a more extended effect of emotion than gaze in the 5 IOC sites of Patient 17. We note that the emotion stimulus exhibits more extensive changes across the face relative to the gaze change, which is confined to the eye region; this was confirmed by the analysis of the degree of changes in the visually presented stimuli (see Supplementary Fig. 2 in (Huijgen et al. 2015)). Additionally, we note that the majority of IOC showed responses to both face onset and social cue change, but a few sites showed responses to one type of face stimulus only, which might be suggestive of different functional regions within our ROI.

### The STC is differentially sensitive to facial motion, and gaze in particular

The most striking finding in this study was the STC particular sensitivity to facial motion (Face 2)—an important finding, due to the relative paucity of available intracerebral neurophysiological data for this region in humans. Specifically, the *gaze change* demonstrated the largest effect sizes in the STS, and indeed overall across all the ROIs (Fig. 5). The active electrodes within the STC ROI typically fell within the mid- to posterior aspect of the STS (MNI y-coordinates ranging from −35 to −53) in both right and left hemispheres. The change of gaze to averted gaze direction relative to the direct gaze condition elicited a larger N200 amplitude and this condition difference could persist beyond N200, out to 400 milliseconds. Importantly, this was accompanied by multiple polarity reversals across adjacent bipolar sites providing evidence of a local generator in STS.

This result is in direct line with the earlier iEEG study by Caruana and colleagues (Caruana et al. 2014) who used monopolar derivations and demonstrated larger N200-like ERPs to viewing gaze aversions relative to direct gaze changes in the STS region. Our study extends these findings by providing direct iEEG evidence for locally generated responses to gaze in the STS on bipolar data. Early scalp EEG potential studies have repeatedly demonstrated larger N200-like potentials to gaze aversions away from the observer relative to transitions to direct gaze for natural face images (Puce et al. 2000, 2003; Latinus et al. 2015; Rossi et al. 2015). Similar MEG changes have also been reported when subjects view ‘interacting’ avatar faces who either looked at each other or to one side of the screen, without at any time looking directly at the observer (Ulloa et al. 2014). Furthermore, a number of fMRI studies (Puce et al. 1998, 2003) indicated that the STS region is sensitive to eye motion, and that averted gaze can produce larger STS activation than viewing direct gaze (Engell and Haxby 2007). Neuropsychological studies of patients with rare— acquired and circumscribed — lesions affecting the superior temporal cortex also showed that impairments in judging gaze direction can occur in these patients (Akiyama, Kato, Muramatsu, Saito, Nakachi, et al. 2006; Akiyama, Kato, Muramatsu, Saito, Umeda, et al. 2006). Intracerebral EEG nicely complements these data from different modalities by providing precise temporal and spatial information. Our findings are among the first to provide direct neurophysiological evidence for STS differential sensitivity to gaze—a particularly interesting finding given the recent postulation of a third visual pathway from STS to V1 (Pitcher and Ungerleider, 2021).

Some interesting questions remain, and should stimulate future research in this area, particularly with respect to functional specialization along STS and also hemispheric differences in STS response properties. For example, a recent 7 Tesla fMRI study using 1 mm^3^ isovoxels investigated the topography of response properties of the human STS to viewing gaze changes, emotional expressions, and speech-related mouth movements in 16 healthy subjects (Schobert et al. 2018). The right STS, in particular, showed a distinct division of labor across its posterior-to-anterior axis: gaze-related activity occurred in its posterior and middle sector, emotional expressions preferentially activated the middle portion of the STS, and the anterior STS was most sensitive to speech-related activity. Although the coordinate limits for the breakdown between these posterior-to-anterior sectors were not explicitly stated, these results appear in agreement with the range of significant gaze effects in our study: bipolar coordinates with a MNI y-range of − 53 to −39 in the right STS and y=−36 and −35 in the left STS (Supplementary Fig. 10). In another study, Deen et al. examined the STS in 3 mm thick slices to a suite of social cognition tasks, including moving face video clips (Deen et al. 2015). The activation along the STS was punctated and culminated at y=−41.1 in the right hemisphere and y=−36.2 in the left hemisphere for the moving faces (Deen et al. 2015). This is again entirely consistent with the neurophysiological data of the current study. It is interesting to note that the coordinates of the bipolar sites seemed somewhat more variable for the emotion effect (with y= −53 to −10 in the right STS and y = −36 and −19 in the left STS —see rightmost bottom inset of Supplementary Fig. 10), as compared to the gaze effect. Future studies will be necessary to fully uncover the functional organization of STS.

Taken together, all of the abovementioned studies across multiple assessment modalities indicate that the STS is critical for monitoring gaze direction and therefore is important for social attention (Puce et al. 2015; Pitcher and Ungerleider 2021).

### Sensitivity to face onset and social cue changes in FC

Consistent responses to face onset and facial social cue changes were observed along the posterior-to-anterior axis of the FC as well as along the STC and in the IOC. Unlike STC, the FC responded vigorously to both stimulus types, with a majority of bipolar sites showing responses to both static face onset and face change. This differential neurophysiological sensitivity concurs with several studies that found activation in the face-responsive fusiform cortex (FFA) to both static and dynamic faces (LaBar et al. 2003; Fox et al. 2009; Pourtois et al. 2010; Pitcher et al. 2014; Pitcher, Ianni, et al. 2019). In particular, in a recent extensive fMRI study in healthy subjects (Pitcher, Ianni, et al. 2019), the fMRI activity in the ventral cortex (FG) did not differ in response to static and dynamic facial stimuli, whereas fMRI activation in lateral temporal cortex (STS) was more strongly driven by dynamic faces and bodies (Pitcher, Ianni, et al. 2019). This is fully consistent with the pattern of responsiveness to face onset and face change that we found in the FC and STC. Moreover, we found both gaze effects and emotion effects on FC sites, with relatively more sites responding to emotion (see Supplementary Fig. 8), although *effect size did not differ for gaze and emotion*. This agrees with the studies that found fusiform activation to dynamic emotional expression (LaBar et al. 2003; Pelphrey et al. 2007). Recently, Bernstein & Yovel (2015) reviewing the literature and taking into account task differences, proposed a model for face processing where they stressed that form and motion are likely to be the primary functional division between a ventral face processing stream (that features the fusiform face area [FFA] and occipital face area [OFA]) and a dorsal stream (that features the STS), respectively (Bernstein and Yovel 2015). Altogether, our results support the idea that FC regions sensitive to faces extract information about both gaze and emotion, because these dynamic social cues contain important configural information for the recognition of individual identities and their gestural idiosyncrasies (O’Toole et al. 2002; Jenkins and Langton 2003; Knappmeyer et al. 2003; Bernstein and Yovel 2015).

Gaze effects and emotion effects were observed across a large time range of our window of analysis, with effects consistently observed in the N200 time range and extending to 400 ms post-change. Yet, on the few sites where an effect of both emotion and gaze were observed, these effects were observed in non-overlapping time windows, and there was also little interaction between emotion and gaze, in the later time range (~300 ms). These data suggest that FC may process emotion and gaze sequentially, with little interaction at the tested latencies.

### ITC belongs to the face processing network

What was somewhat of a surprise was the robust and consistent neurophysiological response from the ITC ROI. The sites responsive to face onset and/or change were observed in posterior to mid-portions of the ITC. The ITC showed responsiveness to both face onset and face change and about half of the ITC sites that responded to face change were sensitive to the gaze or emotion change, with relatively small effect sizes. Perhaps the small effect size in the ITC has made it more challenging to demonstrate activity using other assessment modalities e.g. fMRI, to these stimuli. Yet, reliable activity was originally described in the ITS to facial motion (Puce et al. 1998), although this region appears to show a greater sensitivity to hand (Pelphrey et al. 2005; Thompson et al. 2007) and body (Pelphrey et al. 2005; Atkinson et al. 2012) motion. How non-selective regions may participate in the processing of faces remains an important open question (Haxby et al. 2001). This issue is clearly beyond the scope of the present study, because face selectivity was not tested here. This notwithstanding, our data support the view of ITC as a face-responsive region, participating in the visual processing of face, gaze, and emotion, even if it may not be *selective* of faces *per se*.

### Interaction between emotion and gaze in the 4 ROIs

It is important to underline that our experimental protocol was not symmetric in the way it manipulated emotion and gaze. After the initial neutral face presentation, the faces turned happy or fearful, while gaze turned sideways or remained direct. Besides, as mentioned above, emotion change involves extensive face motion in comparison to gaze change, which is very local and narrow. That said, we found reliable effects of gaze and emotion in the 4 ROIs, with some sites showing both main effects, but very few sites showing a statistically significant interaction between gaze and emotion, in the [0; 400 ms] time window of our analyses (IOC: 1 site, STC: 3 sites, FC: 2 sites, ITC: 1 site). These interactions were observed in late time windows (beyond 300 ms) in IOC and FC, and in both early and late time windows in STC and ITC. The timing of the integration between the different information extracted from faces, such as gaze and emotion, is a long-standing question (see (Graham and Labar 2012) for a review). In one of our earlier MEG studies, we presented dynamic emotional expressions in different social gaze contexts (Ulloa et al. 2014). Interactions between emotion and gaze were complex, showing different timings over posterior (occipito-temporal) and anterior (fronto-temporal) scalp regions. Interestingly, over posterior sensors, emotion effects independent of gaze were initially observed, followed by an interaction between emotion and gaze (Ulloa et al. 2014). Although this study and the present one differed in numerous aspects, they agree in suggesting that visual occipito-temporal regions *can process emotion and gaze independently in the initial stages of face processing*, at least under the circumstances of the experimental protocols used. In the current study design, our epochs were limited to 400 ms, due to the multiple stimulus design, so we were not able to monitor whether or not late interaction effects occurred. Future studies examining these variables will have to use designs that can study and potentially dissociate these interactions by presenting these different components of facial motion at different times.

### A general comment regarding sensitivity to static versus dynamic stimuli

Within the extensive iEEG field potential literature dealing with face processing, most studies have focused mainly on the fusiform gyrus and its patterns of responsivity to static faces, in line with the large existing literature in fMRI also using static faces. This iEEG study and that of Huijgen et al., 2015 (Huijgen et al. 2015) from our lab are the only studies, to our knowledge, where comparisons between intracerebral responses to *these different stimulus types* have been directly performed in the same subjects and experiment. A comparison such as this is crucial for sorting out a valid processing hierarchy and incorporating the results of laboratory-based (static) and naturalistic viewing tasks into ecologically valid models of face processing. Traditionally, the literature has kept these two dimensions separate—largely due to two influential models of (familiar) face processing (Bruce and Young 1986; Haxby et al. 2000). Yet, motion is an essential dimension of a face in everyday life, irrespective of whether the individual’s identity is being sought or whether the meaning of a facial expression or an eye gaze change has to be decoded. The demarcation in the literature between the two face processing pathways has been somewhat academic and artificial—as noted by the assessment of data in both human and non-human primates (Pitcher and Ungerleider, 2021). Accordingly, our data indicate that both ventral and dorsal pathways extract *all types of facial information* (Fairhall and Ishai 2007; Bernstein and Yovel 2015)—at least to a basic level when interacting with a dynamic face. We acknowledge that we have used apparent motion in the current study and not a continuous motion stimulus. However, apparent motion and motion simulation have been shown to generate comparable neurophysiological effects (see (Puce et al. 2003; Ulloa et al. 2014; Latinus et al. 2015)).

### Different routes of facial information flow in cortex

Over the past few decades the predominant view has been that the information flow in the core face processing network takes two main cortical routes, mapped onto the ventral and dorsal visual pathways and processing the invariant and variant aspects of face, respectively (Haxby et al. 2000; Gobbini and Haxby 2007). Yet, as already noted from the existing literature it is still not clear how information is exchanged between the STS and FFA, given the known absence of abundant and direct white matter connections between them (Ethofer et al. 2011; Gschwind et al. 2012; Pyles et al. 2013; Grill-Spector et al. 2017). Additionally, there has been the view that visual processing along the ventral pathway follows a hierarchy, based on fMRI studies where the sluggish hemodynamic response has been observed to occur earlier in posterior structures such as OFA relative to the FFA (see Fig. 10A and 10B; but see Jiang et al 2011). We have already alluded to issues with making assumptions about timing from hemodynamic data. For instance, the blood flow response likely will contain neurophysiological responses across a large timescale e.g. N200s to N700s, as well as oscillatory activity (Puce et al. 1997). Furthermore, the vascular irrigation across various parts of occipito-temporal cortex relies on different major feeder vessels (Marinkovic et al. 1987). When data from other assessment modalities, in addition to fMRI, are considered, there are inconsistencies in the dual visual pathway framework, leading to the proposal of a third visual pathway from V1 via MT/V5 to the STS (Pitcher and Ungerleider 2021). White matter tractography studies have indicated that the IOG is connected to ventral face responsive regions (OFA, FFA and mid-fusiform gyrus) via the ILF and shorter-range occipito-temporal tracts that show more or less overlap with ILF (Catani et al. 2003; Pyles et al. 2013; Grill-Spector et al. 2017). Our data concord partly with this idea. We found some overlap between IOC—and to lesser extent FC—active sites and posterior ILF and additionally a scarce overlap between ITC sites and anterior ILF. Our data are sparse since we had a limited number of active sites, particularly in the most anterior temporal regions. Yet, they may be rather in line with the recent emphasis on the importance of short-range white matter tracts (not included in our analysis) in information flow within the face network (Gomez et al. 2015; Wang et al. 2020).

**Figure 10.**
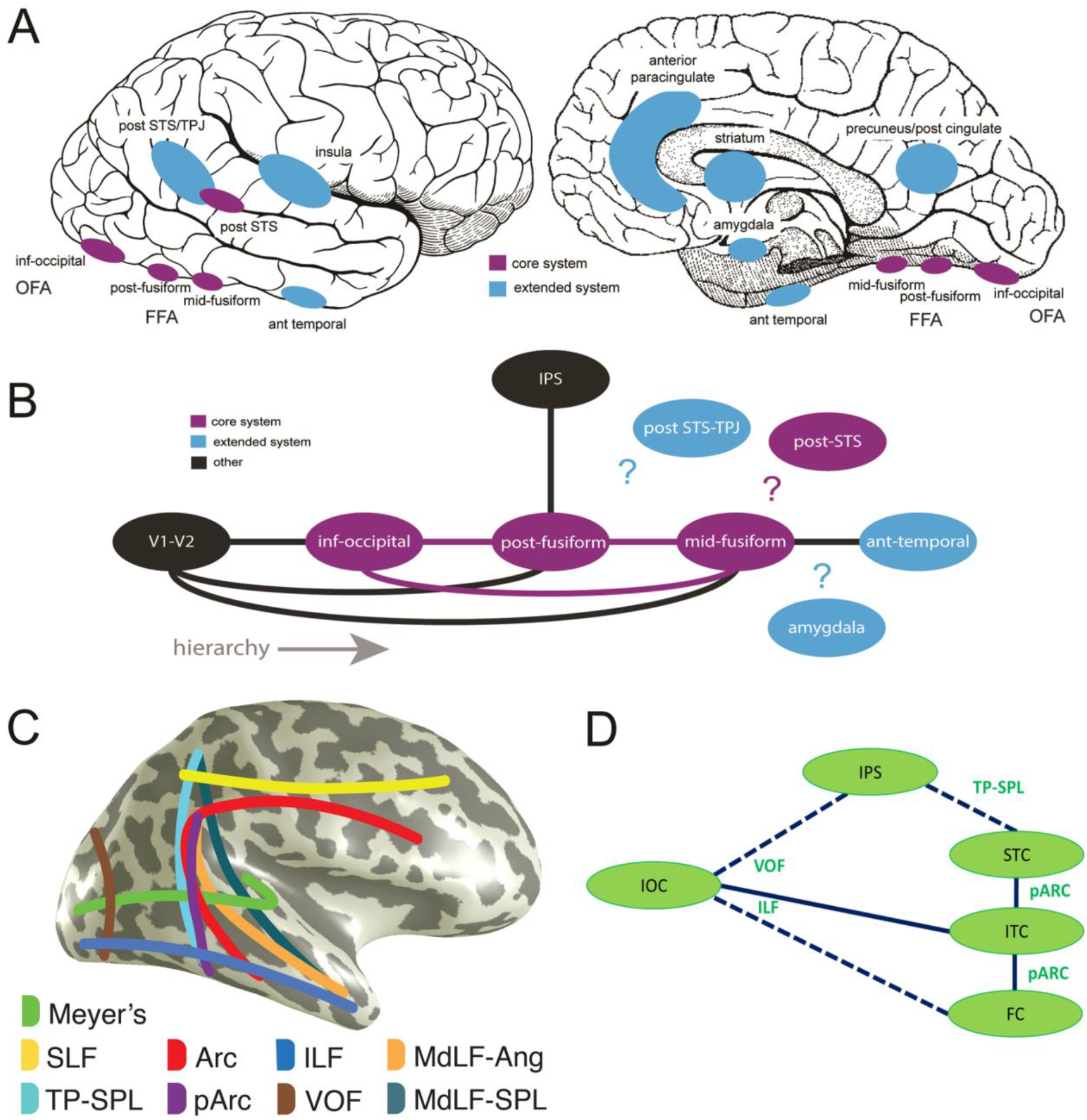
Putative routes of information flow for faces in the brain. **A.** Structures of the core (pink) and extended (blue) face network based on Gobbini & Haxby et al. 2007 and Haxby et al. 2000. Legend: ant = anterior, post = posterior, inf = inferior, OFA = occipital face area, FFA = fusiform face area, STS = superior temporal sulcus, TPJ = temporo-parietal junction. **B.** Putative hierarchy of information flow within the face network showing known direct white matter connections (solid lines) (adapted from Grill-Spector et al., 2017). The nature of connections between other brain regions, including the posterior STS (part of the core face network) and amygdala (extended face network) are not known. Legend: similar to part A, IPS = intraparietal sulcus, V1-V2 = early visual areas. **C.** White matter pathways which may be involved in routing information within the posterior visual face pathway, based on overlap analysis of white matter tract endpoints in 1066 healthy subjects and coordinates of active bipolar sites in patients with epilepsy. (Modified from (Bullock et al. 2019)). Legend: superior longitudinal fasciculus (SLF), temporo-parietal connection of the superior parietal lobule (TP-SPL), posterior arcuate fasciculus (pArc) and arcuate fasciculus (Arc), inferior longitudinal fasciculus (ILF), vertical occipital fasciculus (VOF), and middle longitudinal fasciculus (MdLF) branches of the angular gyrus (Ang) and of the superior parietal lobule (SPL). We also included Meyer’s loop (Meyer’s) that is the optic radiation connecting the lateral geniculate nucleus and the occipital lobe in this schematic figure. **D.** Putative routes of information flow evidenced by the multimodal data integration of neurophysiological data and white matter tract endpoints overlap analysis. The solid lines represent the putative routes for which an overlap between both ends of tract endpoints and active sites was observed, while broken lines indicate connections with overlap at one end of the tract only, in this study. Note that short-range fibers that may be key in information flow across the ventral occipito-temporal ROIs (from IOC to ITC and to FC) and could also play a role in connecting FC, ITC, and STC were not included in our tract endpoint analysis and are not represented here.

We undertook the exploration of likely white matter pathways that might propagate visual information related to social/emotional and facial attributes in an independent group of (healthy) subjects, using the MNI co-ordinate locations of active bipolar sites from the patients as a reference, because of the difference in ERP patterns across the ROIs for Face 1 and Face 2 stimulus types. Indeed, our data pose a challenge for information transfer between IOC and STS as proposed by Fairhall and Ishai (2007), because of the large latency (and morphology) differences between the ERPs of the IOC and STC to face onset, which contrasted with the latency difference between IOC and FC / ITC to face onset and with the ERP pattern to face changes (see Fig. 4D). Thus, ERP latencies suggested a sequence of activation where information related to face onset (Face 1) would travel in parallel with equal speed from IOC, to ITC and FC, and then reach STC after a delay. In the STC, the activity for Face 1 was quite late, broad and slurred relative to the other three regions—begging the question of whether the information might *take a different route* relative to that reaching the FC and ITC. For responses to apparent facial motion (Face 2), the IOC also had the shortest latencies, suggesting that this was again the entry point into the system. However, in this case, responses with approximately equal latencies were observed in FC, ITC, and STC. Altogether, these results suggest that there exist *at least* two routes—one indirect and one more direct or faster—from IOC to STC. Hence, we were interested in looking for the alternative routes for the information to take from IOC to reach the STC.

Overall, our exploratory overlap analysis gave a few clues as to the routes that may subtend the sequence of activation in the system as described above (see Fig. 10D). Our initial hypothesis was that the information might be conveyed to dorsal visual system structures such as parietal cortex (e.g., intraparietal sulcus; (Puce et al. 1996, 1998)) via the VOF (see Fig. 10D). The parietal cortex would then parse information to the STC (specifically the STS) via a more anterior dorsal-ventral white matter route, involving pArc and/or TP-SPL (Bullock et al., 2019). Our data from IPS do not allow us to make this conclusion—our sampling in this region was very sparse and we did not find overlap between the few IPS active sites and superior endpoints of either tract. This idea is, however, plausible given that: (i) previous fMRI studies have documented activation to viewing gaze changes, mouth movements, and even static faces in IPS and STS (Puce et al. 1996, 1998); (ii) there are known white matter connections between the parietal regions and the posterior fusiform, via the VOF—for the most posterior portion of the fusiform— (Yeatman et al. 2014; Takemura et al. 2017; Grill-Spector et al. 2017), pArc (Weiner et al. 2017), and TP-SPL (Bullock et al. 2019). That said, an alternative route may be suggested from our overlap analysis of white matter tract endpoints and coordinates of bipolar active sites in ROIs. The information flow between the ventral stream and STC could proceed from the IOC to the ITC and FC via short range tracts (e.g. U-fibers) and ILF, and then from the latter regions onto the STC via pArc, TP-SPL, or again via some U-fiber tracts. This would constitute an indirect route between IOC and STC, different from the dorsal route mentioned above, and which could account for the ERP dynamics to face onset. Yet it leaves open the question of the more direct or faster route between IOC and STC in response to face changes—which could potentially be the putative third visual pathway via MT/V5. Very recent work comparing visual white matter tract pathways in humans and great apes (Roumazeilles et al. 2020) shows some idiosyncrasies in the ILF (which can be divided into a medial and ‘flat’ component in its posterior end). It is therefore possible that information from V1 could find its way via MT/V5, potentially via the ITC to the STS along the third visual pathway (Pitcher and Ungerleider 2021).

There is currently interest in combining multimodal data to explain the nature of the interactions between the structures in the core and extended face networks (Grill-Spector et al. 2017; Wang and Olson, 2018; Wang et al. 2018, 2020). Studies have used mainly fMRI data— both resting-state and also task-related—to examine functional connectivity differences between the structures in the face network in each cerebral hemisphere (e.g. (Rosenthal et al. 2017)). Notably, left hemispheric effective connectivity analyses suggested a largely feed-forward arrangement from posterior to anterior structures, whereas there was a feed-forward and feed-back flow of information within the right hemisphere (Wang et al. 2020)—pointing to the extensive bottom-up and top-down information flow within the right hemisphere face network. Other interesting implications from this study were that: (1) the right hemisphere functional ‘face connectome’ is highly dependent on face-selectivity in individual voxels; (2) it may be valuable to examine interindividual variability in these face connectome maps. From our perspective, we would advocate that the incorporation of invasive neurophysiological data, with its high temporal resolution, will be crucial for shedding light on the timing of the interactions between the various structures of the face network, and perhaps be able to finally determine who is the ‘cart’ and who is the ‘horse’.

### Limitations and relative strengths of our study

Intracerebral recordings from epileptic patients always present limitations in that the patient population is necessarily different from the usually studied healthy population. The intracerebral electrodes are implanted to identify potential seizure foci - and will necessarily have sites that have been targeted for the identification of epileptic EEG activity. These depth electrodes are typically targeted at the long axis of the hippocampus and the amygdala. To deal with this issue, here we used extensive and strict artifact rejection procedures, to restrict the presence of abnormal activity in the data that were analyzed. This also necessitated limiting the activity we could examine. That said, all patients had appropriate behavioral responses during the task, no impairments in face processing and normal anxiety levels.

Despite adequate behavior, brain anatomy in these patients may not necessarily be neurologically normal. Additionally, tissue distortions introduced by the intracerebral electrodes themselves can further make the anatomy challenging to identify for automated procedures for electrode localization. For these reasons, we proceeded manually, using the individual anatomical scan of each patient (before and after implantation), to identify the precise anatomical structure (gyrus/sulcus/white matter) where each electrode recording site was located. In our opinion, this method led to a much higher anatomical precision. Still, our sampling of occipito-temporal regions was not exhaustive (as shown in Fig. 2), and we did not sample some additional anatomical structures involved in face processing (e.g. the insula; Caruana et al. 2014). In particular, we cannot directly compare the results between the current study and our earlier study with the same protocol (Huijgen et al. 2015). The latter study focused on the ERPs from amygdala contacts, based on 5 patients from the same initial cohort. In our present analysis, however, some patients from Huijgen et al. (2015) were excluded because we scrutinized the entire neurophysiological dataset. Since there was persistent interictal EEG activity in some of the non-amygdala contacts, this resulted in too few trials for our final data analysis of occipito-temporal ROIs. We may just note that for the amygdala data of Huijgen et al. (2015), response latencies varied, but the earliest responses to the *gaze change* in the right amygdala occurred at potentially comparable latencies to those of the cortical responses reported here. Responses to gaze were also seen more clearly relative to responses to emotion, the latter being highly variable. Intracerebral EEG data are rare and rich - however complex—and we think that the extended analysis of the iEEG ERP responses from occipito-temporal regions provide a fruitful complement to our previous amygdala-centred study.

In our study, we did not use localizer tasks to identify functionally defined regions such as MT/V5, EBA, OFA, where a selection of different visual categories would have needed to be explored (e.g., Fig. 9, Puce et al. 1999). This is why we discussed our results above, particularly from the IOC ROI which was likely to encompass the OFA and MT/V5, in relation to the responsiveness to faces but also to biological motion. A limitation of typical functional localizers is however related to the use of static stimuli (e.g., Rangarajan et al. 2020). Such localizers may not have allowed identifying those sites where we observed responses to the face emotion and/or gaze changes but not to face onset here, particularly important in the STC ROI.

Another question here relates to the relative role of the right and left hemispheres in the processing of gaze and emotion. In our patient sample, most of the seizure disorders were predicted to be, and were localized, to the right hemisphere. Because of this implant bias, our sampling of the left hemisphere was quite limited. This notwithstanding, we found similar results across hemispheres. What is not clear is whether there is really no difference between the hemispheres, or this is a consequence of additional recruitment of the left hemisphere because the right hemisphere has been compromised by the seizure disorder.

Intracerebral recordings are not immune to volume conduction effects, which can confound the precise localization of neural generators, as often the reference can be located at considerable distance from the depth electrode contacts (e.g. on the scalp). To circumvent these issues we computed bipolar montages. Moreover, the bipolar sites where our effects of interest were observed showed very focal and clearly observable polarity reversals, on occasions even showing multiple local polarity reversals over a very short distance (e.g. Patient 17, STS activity, see Fig. 7). This indicates that the reported effects are likely localized to these regions.

One could debate the validity of our comparative analyses comparing active sites in patient invasive neurophysiological data with those of white matter pathways in a healthy population of subjects. It may however be noted that good agreement between the activation data of healthy subjects and the neurophysiology of epileptic patients has been previously described (Puce et al. 1995), including in a patient with a seizure-onset-zone in the fusiform gyrus (Puce et al. 1997).

A major strength of our study resides in the analysis of several hundreds of bipolar-sites, pooled together over 11 patients. This does raise issues of multiple comparisons. That said, our data are reliable in the sense that we had a large number of experimental trials per condition and used a cluster-based approach, rigorously corrected for multiple comparisons within each of the 4 ROIs studied. We went beyond null hypothesis significance testing, by complementing our data analysis with effect sizes, to evaluate the robustness of our effects. Our paradigm also allows us to study the effects of gaze and emotion in the same recording epoch.

## CONCLUSIONS

In this study, we examined intracerebral responses to face onset and facial social cue change (gaze, emotion) in the inferior occipital, fusiform, inferior temporal, and superior temporal cortices. We found robust responses to the different stimulus types in the four ROIs, supporting the view that various facial attributes are processed in parallel in occipitotemporal cortex. However, certain stimulus dimensions, namely gaze changes, preferentially activated the STC relative to other brain regions. The IOC appeared as a likely common entry point into the ventral (FC, ITC) and dorsal (STC) face processing system, through the inferior longitudinal fasciculus and the vertical occipital fasciculus. Communication across the face responsive regions also involves the arcuate fasciculus and the temporo-parietal connection, providing some potential routes among ventral and dorsal regions. Further studies in a larger set of patients with more abundant intracranial sampling throughout occipitotemporal cortex and combined structural connectivity imaging will have to be performed to map out the activation sequence and route taken for different dimensions of information relating to the face. This is particularly important given that complexity increased with the recent postulation of a third potential route of visual information flow in the brain.

## ACKNOWLEDGMENTS

The Authors thank Fanny Lachat and Josefien Huijgen for their contribution in data acquisition. The initial electrode localizations post-implant were performed by Dr Dominique Hasboun, neurosurgeons Pr Stéphane Clemenceau and Dr Bertrand Mathon (AP-HP, GH La Pitié -Salpêtrière-Charles Foix, Neurosurgery Department, F-75013, Paris, France) implanted the electrodes. We are also grateful to Pr Vincent Navarro (head of the Epileptology Unit) and to the Staff from the Epilepsy Service who made accommodations for the research studies, and to the patients who willingly gave their time to participate in the study.

## NOTES

### Author contributions

MBR, AP, and NG designed the analysis of the neurophysiological study. MBR and AP executed the analysis of the neurophysiological study with assistance from VD. NG, AP, DB, and FP designed the anatomical connectivity analysis, which was executed by DB. VD and MBR participated in neurophysiological data acquisition under the supervision of CA. LH and KL brought technical support to data acquisition and processing. VD, VL, and CA performed the clinical assessment of the patients. NG designed the original neurophysiological study, which was adapted to epilepsy patient in collaboration with VD. MBR, AP, and NG wrote the manuscript with revisions from the other authors.

