## Supplementary Material for "Visual information routes in the posterior dorsal and ventral face network studied with intracranial neurophysiology, and white matter tract endpoints"

### S1. SUPPLEMENTARY METHODS

#### *S1.1. Labelling of bipolar sites*

Monopolar contacts came to lie either in grey or white matter within a given anatomical structure. Hence, the anatomical locations of the bipolar sites were labelled according to the three following criteria:

1) if the labels of the monopolar contacts were identical (grey/white matter, anatomical structure), then the bipolar site kept the same label as the monopolar contacts;

2) if one of the monopolar contacts was labelled as white matter, the resulting bipolar site took the label of the remaining grey matter monopolar contact;

3) if the two monopolar contacts were in grey matter but with different anatomical labels, we went back to the native post-implantation MRI to determine visually which grey matter structure was closer to the midpoint between the two monopolar contacts, in order to define the anatomical label as precisely as possible.

#### *S1.2. Cluster-based permutation t-test*

In order to test if the activity at bipolar sites significantly differed from zero, we performed a cluster-based permutation t-test at each site (Maris and Oostenveld 2007). We computed a two-tailed t-test against zero across trials at each time point of the z-scored bipolar data epochs (between 0 and 400 ms relative to relevant stimulus onset) and we clustered together the adjacent time points for which the p-value was below 0.01. The obtained clusters were characterized by the sum of the t-values of their individual data points. We then computed the probability distribution of the clusters under the null hypothesis. For this, we randomly permuted two labels across trials: “real iEEG activity”, and “zero”, where “zero” was the label for the vector of zeros used in the initial t-test. We computed the t-test time point by time point as before, clustered the statistically significant data points, computed their sum of t-values and extracted both the maximal positive and minimal negative sums of t-values obtained across the clusters. We repeated this procedure 10,000 times, so as to estimate the permutation distribution of the maximal and minimal sum of t-values statistics (that is, the distribution of this cluster-based test value under the null hypothesis). We then computed the Monte-Carlo simulation p-value of the cluster-level test statistics of the original data as the proportion of test values in the permuted distribution of maximal (or minimal) cluster-level statistics which exceeded (or was inferior to) the originally observed cluster-level test statistics.

### S2. SUPPLEMENTARY RESULTS

#### *S2.1. Behavioral data*

***Supplementary Table 1:*** ***Behavioral data and number of analyzed trials in individual patients.*** *Number of completed blocks, reaction times, hits, false alarms for detecting targets, as well as final number of trials after artifact rejection are presented. SEM=standard error of the mean.*

| Patient | # Blocks completed | Reaction times (mean ± SEM, in ms) | Hits (%) | False alarms (%) | Final number of trials |
|---|---|---|---|---|---|
| 1 | 8 | 541 ± 4.9 | 99.6 | 0 | 779 |
| 2 | 3 | 375 ± 4.7 | 100 | 5.6 | 234 |
| 3 | 8 | 325 ± 4.0 | 93.2 | 9.4 | 777 |
| 8 | 8 | 349 ± 2.3 | 99.3 | 0 | 632 |
| 10 | 8 | 332 ± 2.5 | 99.3 | 0 | 461 |
| 12 | 8 | 326 ± 3.4 | 99.1 | 7.3 | 511 |
| 14 | 6 | 372 ± 3.3 | 99.7 | 4.2 | 455 |
| 15 | 8 | 352 ± 2.2 | 99.9 | 0 | 354 |
| 16 | 8 | 350 ± 2.0 | 99.6 | 0 | 681 |
| 17 | 8 | 333 ± 3.0 | 99.9 | 1.0 | 635 |
| 18 | 6 | 268 ± 2.8 | 100 | 6.9 | 339 |

*S2.2. Responsive contacts*

***Supplementary Table 2:*** ***Responsive contacts in the four ROIs and in the IPS, in left and right hemispheres.*** *Note that only contacts responding to Face 2 were tested for emotion, gaze and interaction effects.*

| ROI | Patient | # contacts total | # contacts responsive to | | | | | |
|---|---|---|---|---|---|---|---|---|
| | | | Face 1 | Face 2 | Face 1 & 2 | Emotion | Gaze | Interaction |
| **IOC** | 1 | 3 | 0 | 0 | 3 | 0 | 0 | 0 |
| | 2 | 2 | 0 | 1 | 1 | 0 | 1 | 0 |
| | 3 | 3 | 2 | 0 | 1 | 1 | 1 | 0 |
| | 8 | 1 | 0 | 1 | 0 | 1 | 0 | 0 |
| | 10 | 8 | 1 | 1 | 6 | 1 | 0 | 0 |
| | 12 | 5 | 0 | 0 | 5 | 3 | 0 | 0 |
| | 14 | 0 | 0 | 0 | 0 | 0 | 0 | 0 |
| | 15 | 4 | 1 | 0 | 0 | 0 | 0 | 0 |
| | 16 | 0 | 0 | 0 | 0 | 0 | 0 | 0 |
| | 17 | 6 | 0 | 0 | 6 | 5 | 5 | 1 |
| | 18 | 4 | 0 | 2 | 1 | 1 | 0 | 0 |
| | **Total** | **36** | **4** | **5** | **23** | **12** | **7** | **1** |
| **FC** | 1 | 18 | 2 | 6 | 4 | 2 | 2 | 0 |
| | 2 | 4 | 0 | 2 | 1 | 1 | 0 | 0 |
| | 3 | 11 | 4 | 0 | 7 | 0 | 2 | 0 |
| | 8 | 10 | 1 | 1 | 4 | 2 | 0 | 1 |
| | 10 | 6 | 0 | 1 | 5 | 1 | 0 | 0 |
| | 12 | 6 | 2 | 0 | 4 | 0 | 0 | 0 |
| | 14 | 16 | 3 | 1 | 9 | 4 | 0 | 0 |
| | 15 | 0 | 0 | 0 | 0 | 0 | 0 | 0 |
| | 16 | 31 | 4 | 3 | 9 | 5 | 3 | 0 |
| | 17 | 10 | 1 | 2 | 6 | 6 | 2 | 1 |
| | 18 | 8 | 2 | 0 | 0 | 0 | 0 | 0 |
| | **Total** | **120** | **19** | **16** | **49** | **21** | **9** | **2** |
| **ITC** | 1 | 6 | 0 | 1 | 2 | 1 | 0 | 0 |
| | 2 | 1 | 0 | 0 | 0 | 0 | 0 | 0 |
| | 3 | 8 | 1 | 0 | 4 | 2 | 0 | 0 |
| | 8 | 4 | 1 | 0 | 0 | 0 | 0 | 0 |

| | | | | | | | | |
|---|---|---|---|---|---|---|---|---|
| | 10 | 5 | 0 | 2 | 0 | 0 | 0 | 0 |
| | 12 | 4 | 0 | 1 | 2 | 1 | 1 | 0 |
| | 14 | 8 | 3 | 1 | 2 | 0 | 1 | 0 |
| | 15 | 3 | 0 | 2 | 0 | 0 | 0 | 0 |
| | 16 | 10 | 2 | 0 | 0 | 0 | 0 | 0 |
| | 17 | 10 | 1 | 2 | 3 | 2 | 1 | 1 |
| | 18 | 6 | 0 | 1 | 1 | 1 | 0 | 0 |
| | **Total** | **65** | **8** | **10** | **14** | **7** | **3** | **1** |
| **STC** | 1 | 11 | 0 | 0 | 0 | 0 | 0 | 0 |
| | 2 | 11 | 0 | 1 | 3 | 0 | 0 | 0 |
| | 3 | 0 | 0 | 0 | 0 | 0 | 0 | 0 |
| | 8 | 14 | 0 | 4 | 1 | 1 | 1 | 0 |
| | 10 | 12 | 3 | 2 | 4 | 1 | 0 | 1 |
| | 12 | 6 | 0 | 2 | 0 | 0 | 0 | 0 |
| | 14 | 4 | 0 | 0 | 2 | 0 | 0 | 0 |
| | 15 | 9 | 0 | 0 | 1 | 0 | 0 | 0 |
| | 16 | 17 | 0 | 1 | 3 | 1 | 2 | 1 |
| | 17 | 8 | 0 | 3 | 2 | 2 | 4 | 0 |
| | 18 | 10 | 1 | 4 | 0 | 1 | 2 | 1 |
| | **Total** | **102** | **4** | **17** | **16** | **6** | **9** | **3** |
| **IPS** | 1 | 0 | 0 | 0 | 0 | - | - | - |
| | 2 | 0 | 0 | 0 | 0 | - | - | - |
| | 3 | 0 | 0 | 0 | 0 | - | - | - |
| | 8 | 0 | 0 | 0 | 0 | - | - | - |
| | 10 | 2 | 1 | 0 | 1 | - | - | - |
| | 12 | 5 | 3 | 0 | 1 | - | - | - |
| | 14 | 0 | 0 | 0 | 0 | - | - | - |
| | 15 | 13 | 4 | 0 | 1 | - | - | - |
| | 16 | 0 | 0 | 0 | 0 | - | - | - |
| | 17 | 2 | 0 | 0 | 1 | - | - | - |
| | 18 | 0 | 0 | 0 | 0 | - | - | - |
| | **Total** | **22** | **8** | **0** | **4** | **-** | **-** | **-** |

*S2.3. Analysis of responsiveness profiles*

To test for differences in proportions of responsive sites across ROIs, we applied the Fisher's exact test first to the number of responsive and non-responsive sites obtained in each ROI and second to the number of sites responsive to Face 1, Face 2, or both in each ROI. The test was implemented using the package "*rcompanion*" in R (Mangiafico 2015) with the function "*fisher.test*". This test is adapted for small sample sizes as tested here. The final two-sided p-value was based on 10,000 Monte-Carlo randomizations. Post-hoc tests to compare responsiveness between ROIs taken 2-by-2 were performed using the function "*pairwiseNominalIndependence*" and the corresponding p-values were corrected for multiple comparisons by false-discovery rate (FDR) correction.

In our pooled data across right and left hemispheres, the overall responsiveness profile showed some idiosyncratic differences among ROIs (Fisher's exact test on number of non-responsive and responsive sites per ROI, with left and right hemispheres pooled together: $p<10^{-4}$; all 2-by-2 post-hoc comparisons: $p<0.05$, FDR corrected for multiple comparisons; see Figure 3 and Supplementary Table 2). The IOC was the most responsive region with an overall responsiveness rate of 89% of all sites tested; all right hemisphere IOC sites (n=22 out of 22) and most left hemisphere IOC sites (n=10 out of 14) were responsive to Face 1, Face 2, or both. A total of 70% of FC sites (right hemisphere: n=60/80; left hemisphere: n=24/40) and 49% of ITC sites (right hemisphere: n=26/50; left hemisphere: n=6/15) were responsive to the face stimuli. The STC showed the smallest proportion of responsive sites with 36% of its sites (right hemisphere: n=29/68; left hemisphere: n=8/34) found to be responsive to Face 1, Face 2, or both.

In contrast to overall responsiveness, the pattern across ROIs changed when considering responses to Face 1 only, Face 2 only, or both Face 1 and 2 (tested using Fisher's exact test: $p=0.024$; Fig. 3A). In this instance, the STC tended to show a selectivity for Face 2 relative to other ROIs, with responses in 46% of its sites (2-by-2 post-hoc comparisons: STC vs. FC: $p=0.05$, STC vs. IOC: $p=0.06$, FDR corrected for multiple comparisons across the 6 post-hoc comparisons computed between ROIs). In contrast, the IOC and FC behaved similarly (IOC vs. FC: $p=0.41$), with most of the sites responding to both Face 1 and Face 2, and few sites responding selectively to either Face 1 or Face 2. The ITC did not differ from any of the other ROIs with respect to its responsiveness profile (ITC vs. FC: $p=0.34$; ITC vs. IOC: $p=0.41$; ITC vs. STC: $p=0.34$); it showed equivalent proportions of sites responding to each stimulus type (Face 1, Face 2, or both).

*S2.4. Responsiveness to faces of the Intraparietal Sulcus (IPS)*

We also assessed activity in response to faces in the IPS from the 4 patients where sites in this region could be analyzed (n=22 sites; Supplementary Table 2). IPS belongs to the extended rather than core face network, but its vicinity to posterior STC and to white matter tracts of the occipitotemporal region suggested a potential role as an indirect route between the STS and the ventral structures in the face network and led us to consider it in our analyses. Only a few IPS sites show responsiveness to Face 1 (n=12) and/or to Face 2 (n=4), with variable ERP waveforms (see Supplementary Fig. 1 of ERP waveforms in IPS individual sites). Therefore, we did not analyze these sites further and only retain their coordinates for our structural connectivity analysis (see section 3 of Results in the main text).

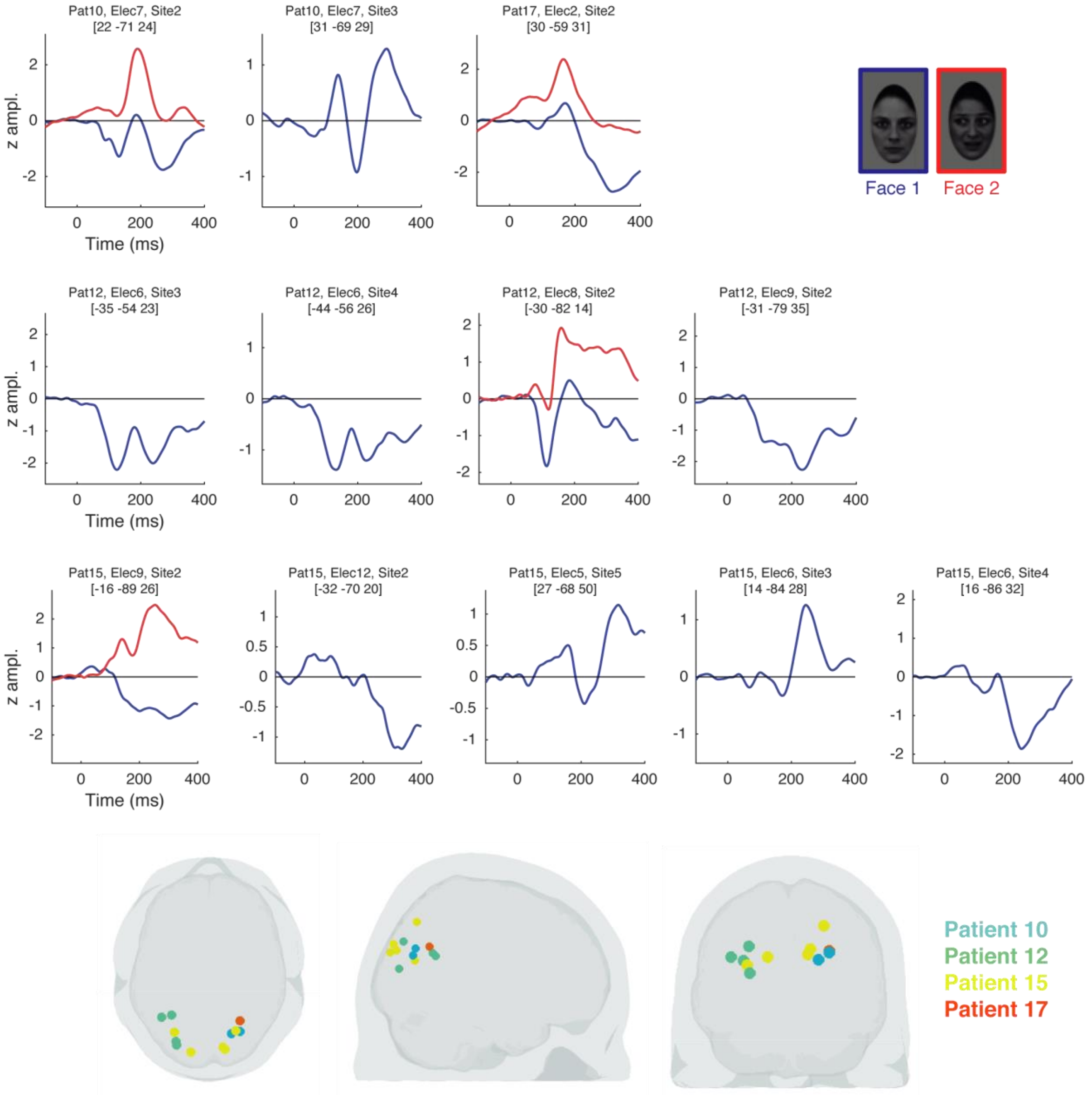

***Supplementary Figure 1:*** ***Individual ERPs to Face 1 and Face 2 from bipolar sites in the Intraparietal Sulcus.*** *These ERPs were obtained with the same methodology and selection criteria as for the four main ROIs. The patient, electrode, site number and the MNI coordinates are indicated for each ERP, whose amplitudes are depicted in terms of z-scores (with no polarity rectification). The responsive site locations can be viewed on the 3D brain silhouette views and the corresponding patient number is identifiable with the dot color (see color-patient correspondence on the right).*

*S2.5. ERP morphology: Individual data*

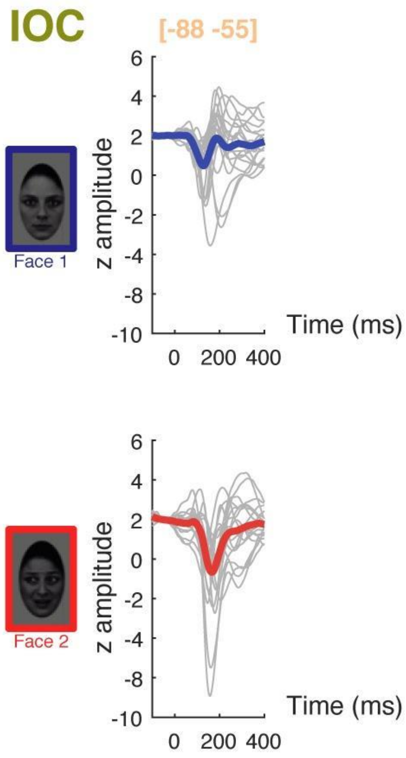


***<u>Supplementary Figure 2:</u> ERPs to Face 1 and Face 2 in the Inferior Occipital Cortex (IOC).*** *The individual data are shown as grey traces. Note that single site ERPs were rectified following the procedure described in Method for the purpose of waveform morphology visualization. The thicker traces correspond to the average (as in Figure 4A).*

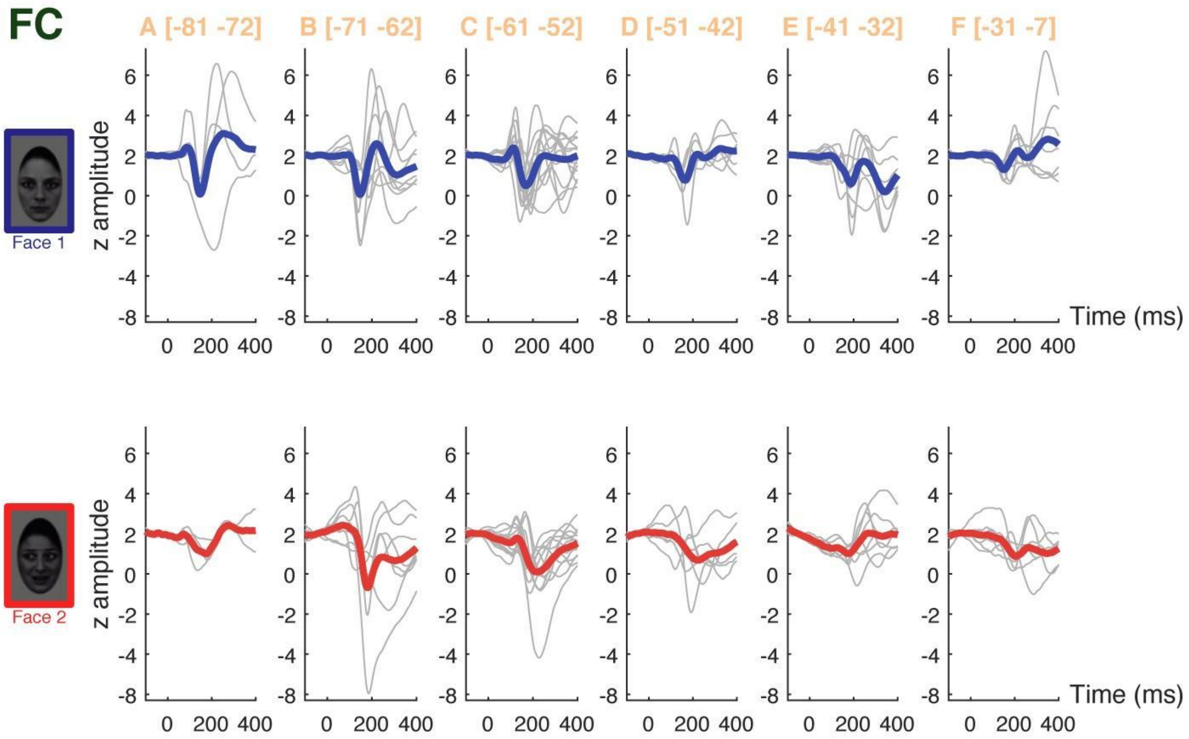


***Supplementary Figure 3:*** ***ERPs to Face 1 and Face 2 in the Fusiform Cortex (FC).*** *Same legend as Supplementary Figure 2.*

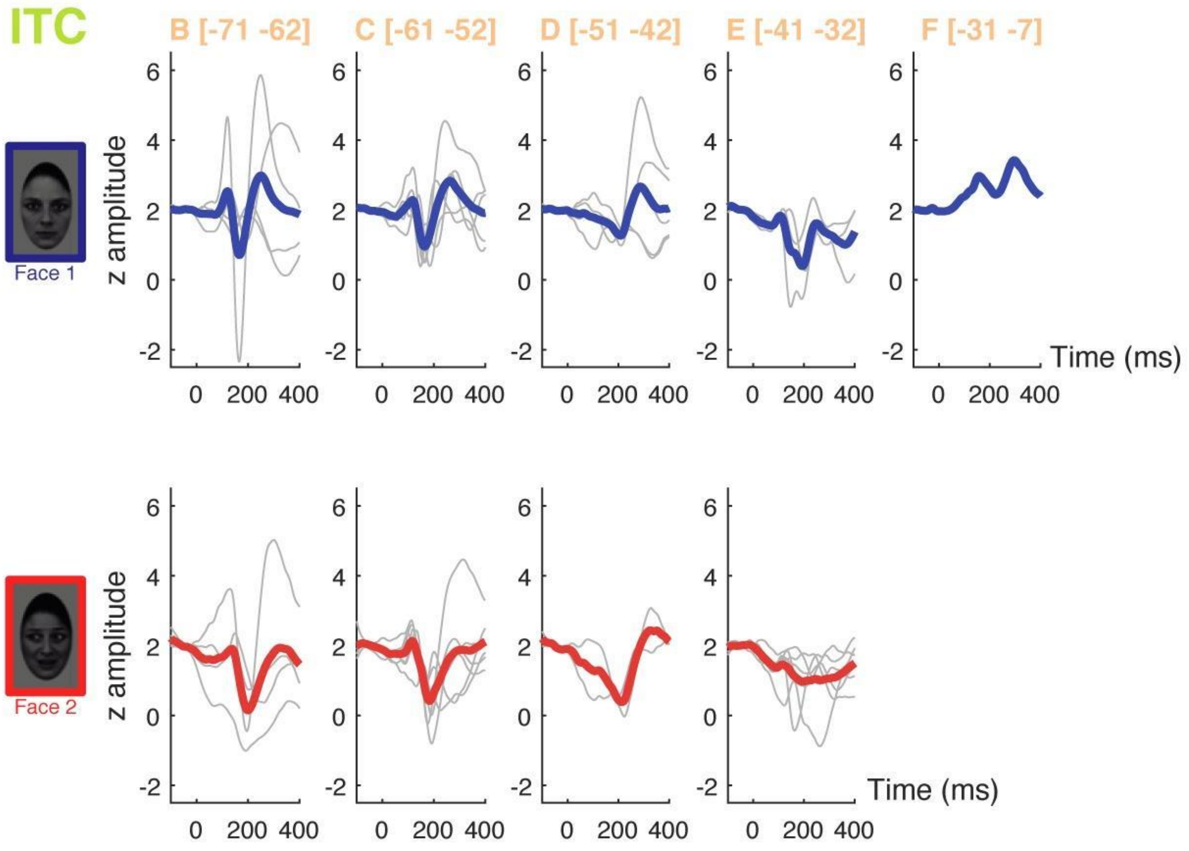


***<u>Supplementary Figure 4:</u>* ERPs to Face 1 and Face 2 in the Inferior Temporal Cortex (ITC).** *Same legend as Supplementary Figure 2.*

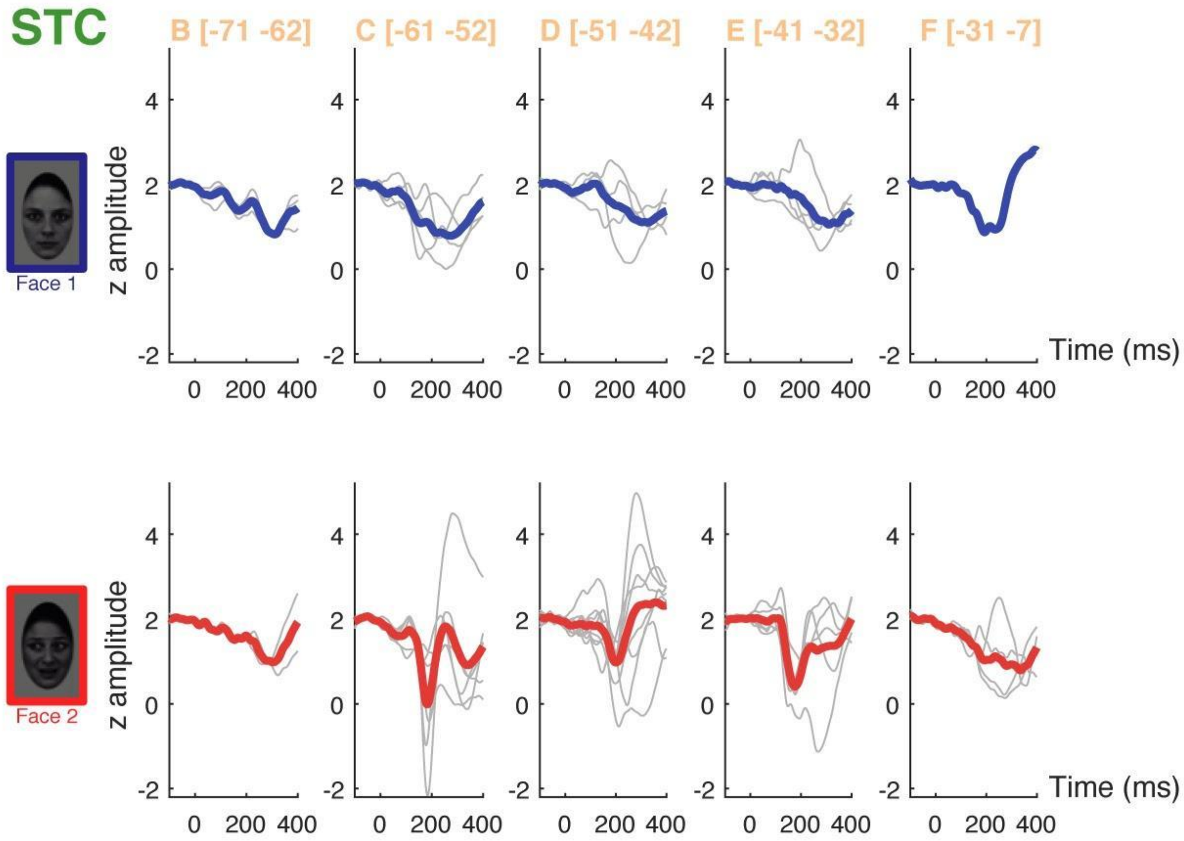


***<u>Supplementary Figure 5:</u> ERPs to Face 1 and Face 2 in the Superior Temporal Cortex (STC).*** *Same legend as Supplementary Figure 2.*

*S2.6. Waveform amplitude and latency: linear mixed effects model analyses*

We here complement the results presented in the part 1.2 of the main text, by testing the effects on ERP amplitude and latency while controlling for inter-patient and inter-contact variability using linear mixed effects models. We computed linear mixed effects models, in R, using the “lmer” function of the package “lme4” (REF Bates et al 2014) and the “anova” function to derive corresponding statistics (Type III Anova table with Satterthwaite’s method).

*ERP amplitude.* We computed a linear mixed effects model to analyse Cohen’s d effect size (absolute value) using the following predictors as fixed effects: Condition (Face 1 or Face 2), ROI (with 4 levels: IOC, FC, ITC, STC), y coordinates (mean centered), the interaction between Condition and ROI and the interaction between ROI and y coordinates. We introduced Patients and Contacts as grouping factors, by including them as random intercepts. There was a significant effect of Condition ($F(1,122.78)=13.76$, $p=0.00031$). The interaction between Condition and ROI was marginally significant ($F(3,120.55)=2.44$, $p=0.067$). Post-hoc tests showed that Face 1 and Face 2 effect sizes significantly differed for IOC ($F(1,33.37)=5.59$, $p=0.024$) and ITC ($F(1,23)=16.92$, $p=0.00042$), but not for FC ($F(1,73.06)=2.19$, $p=0.14$) nor STC ($F(1,23.84)=0.17$, $p=0.68$). These results are consistent with our main findings.

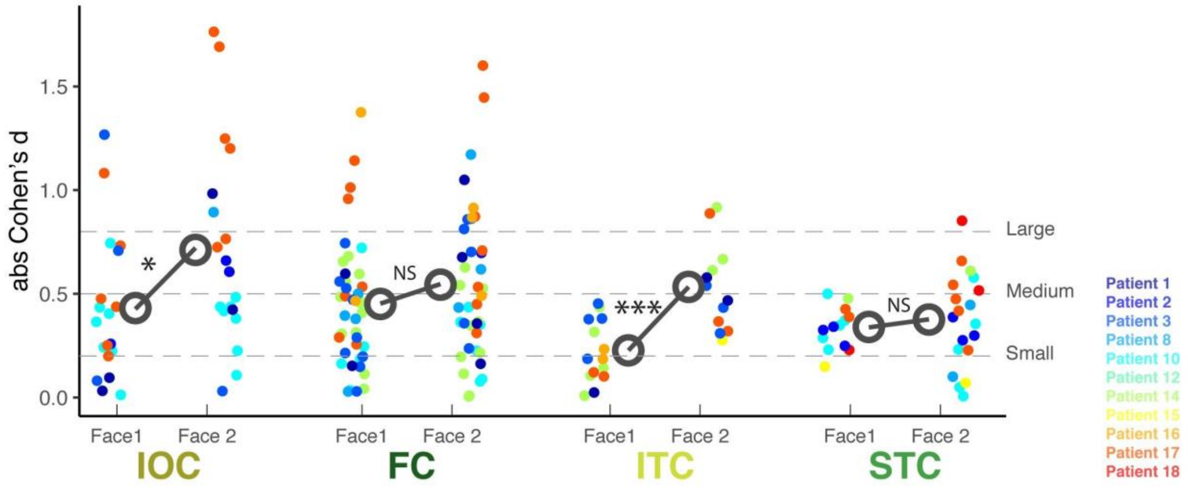


***<u>Supplementary Figure 6:</u> Comparison of ERP effect sizes for Face 1 and Face 2, for each of the ROIs.*** *The dots represent the absolute effect size observed for every contact included in the analysis, color-coded per patient. The big open circles represent the average, for each ROI and face stimulus. NS: non-significant; *: p<0.05; ***: p<0.001.*

*Latencies.* The jackknife-computed ERP latencies used in our main analysis are non-independent; they are therefore not compatible with linear mixed effects models. Thus, for this latter analysis, we needed to extract the ERP peak latencies from each site. For this, we first defined windows of measurement for each ROI and condition: we computed the mean ERP across contacts and patients within each slice of interest and measured the ERP peak latency on this average; we then defined the lowest bound of the measurement time window as the earliest ERP peak latency obtained across slices minus 25ms and the highest bound as the latest ERP peak latency plus 25ms. The minimum ERP latency was then picked in these windows of measurement for each site, in each ROI and condition.

We first tested the effects of Condition and ROI, by computing a linear mixed effects model with the following fixed-effect factors: Condition (Face 1 or Face 2), ROI (4 levels), y-coordinate (mean centered), the interaction between Condition and ROI, the interaction between ROI and y-coordinate. Patient and Contact numbers were entered as random intercepts. We found a significant effect of ROI ($F(3,79.24)=16.54$, $p<0.0001$) and of the interaction between Condition and ROI ($F(3,97.69)=79.11$, $p<0.0001$) on ERP latencies, but no main effect of Condition ($F(1,99.92)=1.66$, $p=0.20$).

As in the main analyses (see main text, paragraph 1.2), we then tested for differences in latency between each pair of ROIs, for each condition. For this, we used linear mixed effects models restricted to the condition of interest, with ROI (2 levels) and y-coordinate as fixed effects and Patient as random intercept. We corrected for multiple comparisons using Bonferroni correction (that is, p values were multiplied by 6, corresponding to the 6 2-by-2 tests performed). The results were consistent with the main analyses (see Suppl Fig 7).

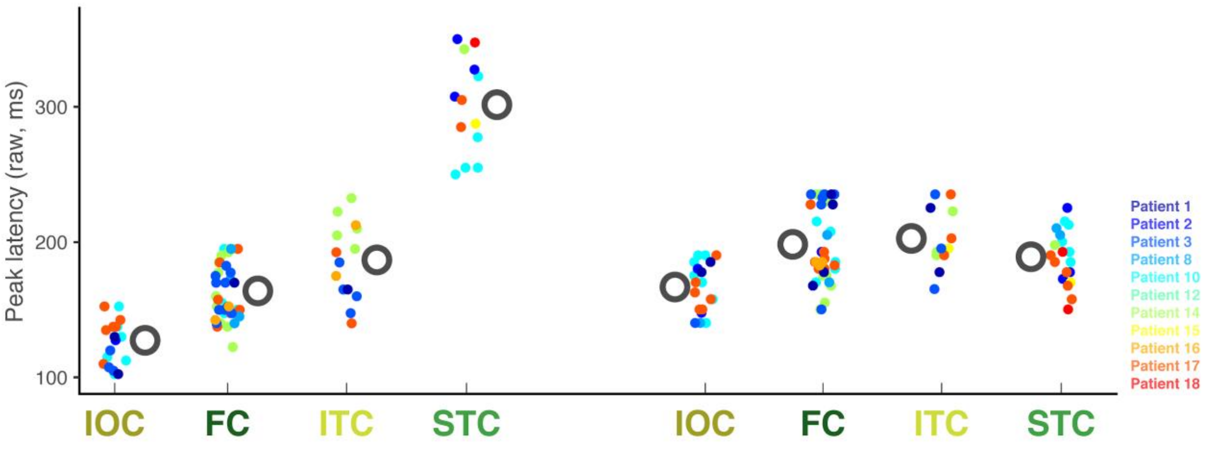


| | FC | ITC | STC |
|---|---|---|---|
| IOC | *** | * | *** |
| FC | | * | *** |
| ITC | | | *** |

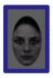
Face 1

| | FC | ITC | STC |
|---|---|---|---|
| IOC | * | (*) | NS |
| FC | | NS | NS |
| ITC | | | NS |

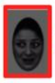
Face 2

***<u>Supplementary Figure 7:</u>*** ***Comparison of ERP latencies for each of the ROIs, for Face 1 and Face 2.*** *The dots represent the peak latency measured on each of the contacts included in the analysis, color-coded per patient. The big open grey circles represent the average peak latency, for each ROI and face stimulus. The tables underneath the plot show the statistical differences between latencies in each ROI taken two-by-two as obtained from linear mixed effects models. NS: non-significant; (*): p<0.06; *: p<0.05, ***: p<0.001 (Bonferroni-corrected).*

Furthermore, we performed two additional linear mixed effects models to test the sequential order of peak latencies which was suggested by our main analyses for Face 1 and Face 2, respectively. For Face 1, we tested the following sequence of latencies: IOC, followed by FC and ITC, in turn followed by STC. For this, we coded the ROI factor as a numerical variable, setting 1 for IOC, 2 for FC and ITC, and 3 for STC. The model also included y-coordinate as a fixed effect and Patients and Contacts as random intercepts. The effect of the numerical ROI factor was significant, confirming the sequence of peak latencies across our 4 ROIs ($F(1,78.97)=96.58$, $p<0.0001$). For Face 2, we tested the hypothesis of earliest latency in IOC, followed by the three other ROIs. For this, we coded the ROI factor as a numerical variable, setting 1 for IOC and 2 for the other ROIs. As in the previous model, we also included y- coordinate as a fixed effect and Patients and Contacts as random intercepts. We found that the effect of the numerical ROI factor was significant ($F(1,86.91)=14.81$, $p=0.00023$), confirming that ERPs in IOC were again the earliest ones.

*S2.7. Sensitivity profiles to social cues across ROIs*

The proportion of sites where statistically significant effects of emotion and/or gaze were observed within each ROI was analyzed using the Fisher's exact test with *fisher.test* function from the package "*rcompanion*" in R (Mangiafico 2015). First, we compared the number of sensitive sites (viz. sites showing some statistically significant effect of emotion, gaze, or interaction between emotion and gaze) and non-sensitive sites across ROIs and second, we compared the number of sites showing an effect of emotion, gaze, or an interaction between emotion and gaze across ROIs. The final two-sided p-values were based on 10,000 Monte-Carlo randomizations.

Overall, between 45% and 71% of the sites responsive to Face 2 showed a statistically significant effect of social cues within each ROI (Supplementary Table 2; number of sites responding to emotion, gaze, and/or showing an interaction, over the total number of sites responding to Face 2: IOC: n=20/28, FC: n=32/65, ITC: n=11/24, STC: n=18/33). The proportion of sites sensitive to social cues did not differ significantly among the 4 ROIs (Fisher's exact test on numbers of sensitive and non-sensitive sites per ROI, pooling together left and right hemispheres: p=0.63). The number of sites showing an effect of emotion, of gaze, or an interaction between gaze and emotion did not differ across ROIs either (Fisher's exact test: p=0.39) (Supplementary Table 2). Emotion or gaze main effects were the most frequent, and only a small number of sites showed an interaction effect. Sensitive sites were observed in both right and left hemispheres (Supplementary Fig. 8-11).

*S2.8. Comparative sensitivity to social cues: linear mixed effects model analyses*

To further support the results of gaze and emotion sensitivity across ROIs, we performed additional linear mixed effects model analyses. This method allows us to take into account inter-patient variability. Note that these analyses constitute alternatives to the approach undertaken in the main text and were performed for purely confirmatory purposes.

We first computed a linear mixed effects model to analyse the effect sizes (absolute values), with social cue (emotion or gaze), ROI, the interaction between social cue and ROI, and hemisphere as fixed-effect predictors, and Patient as a random intercept. This analysis showed a main effect of ROI ($F(3,64.38)=3.26$, $p=0.027$) and of Condition ($F(1,59.82)=4.85$, $p=0.032$) and more interestingly a significant interaction between ROI and Condition ($F(3,61.81)=6.72$, $p=0.00054$). This supports our main results, showing that the ROIs responded differently to the different social cues.

We then restricted our analysis to the effect of gaze in order to test the effect size for gaze across ROIs. The linear mixed effects model on gaze effect sizes (with ROI and Hemisphere as fixed effects and Patient as random intercept) confirmed that gaze effect sizes significantly differed across ROIs ($F(3,23)=6.14$, Bonferroni corrected $p=0.0064$). In contrast, when considering effect sizes for emotion (with a linear mixed effects model on emotion effect sizes), we found no significant effect of ROI ($F(3,41)=2.10$, Bonferroni corrected $p=0.23$). Finally, following the same logic as in the main text, we tested the difference between gaze and emotion effect sizes for each ROI, with linear mixed effects models including Condition as fixed effect and Patient as random intercept. Consistently with our main results, effect sizes for gaze did not significantly differ from effect sizes for emotion in either IOC ($F(1,17)=0.56$, $p=0.46$), FC ($F(1,27.94)=0.30$, $p=0.59$), or ITC ($F(1,6.65)=4.81$, $p=0.066$). They showed a marked difference in STC ($F(1,13)=9.22$, $p=0.0096$).

*S2.9. ERPs in response to social cues for each ROI*

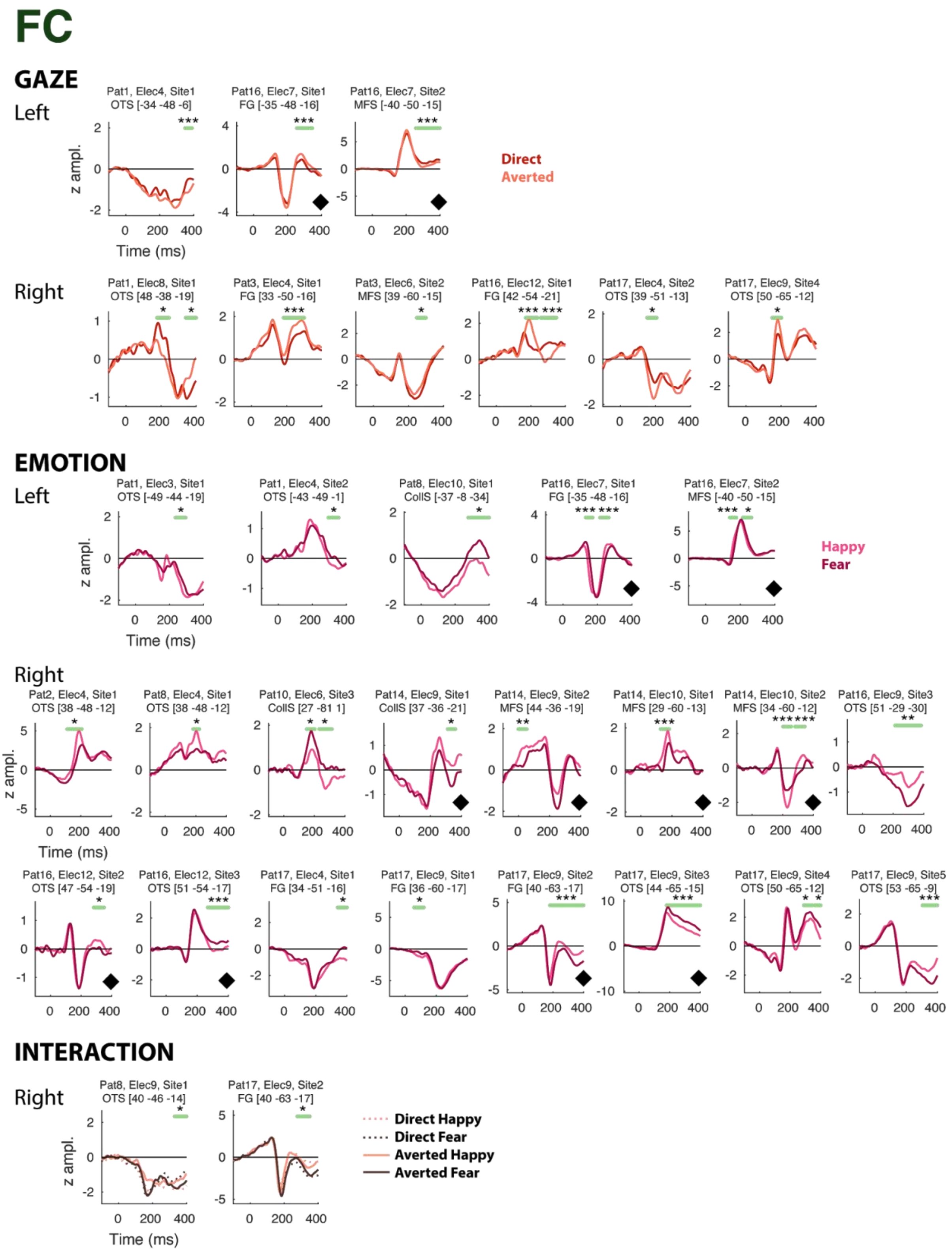


***Supplementary Figure 8:*** ***ERPs from sites in the Fusiform Cortex region (FC) responding to social cues.*** *The patient, electrode and site number, as well as the anatomical localization and the MNI coordinates are indicated for each ERP. Horizontal green bars indicate the time window where the conditions significantly differ. Black diamonds near the x-axis of the plots indicate polarity inversions in adjacent sites. Legend: CollS: Collateral Sulcus, FG: Fusiform Gyrus, MFS: Mid-Fusiform Sulcus, OTS: Occipito-Temporal Sulcus. *: p<0.05, **: p<0.01, ***: p<0.005, Monte-Carlo p-values.*

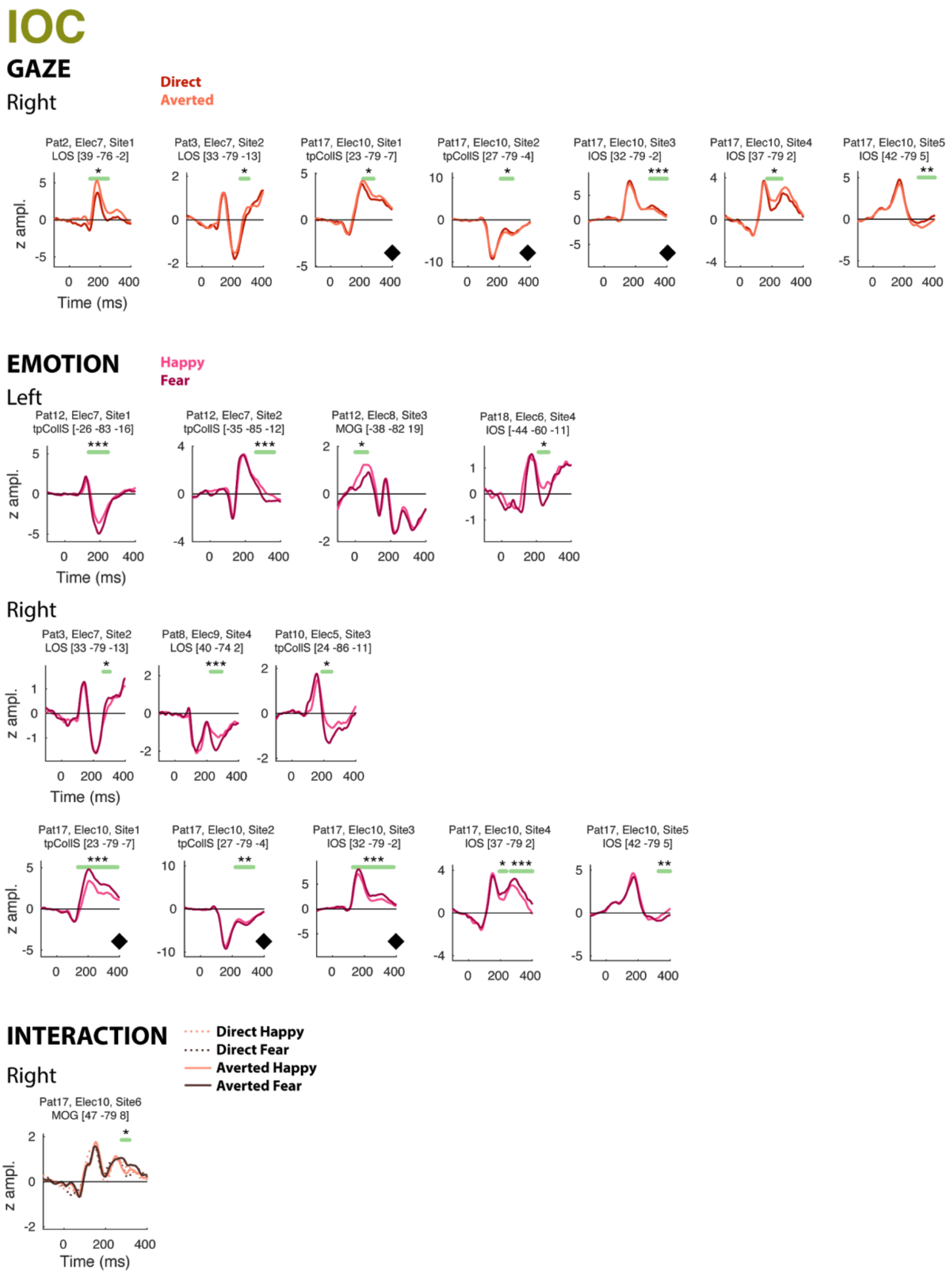


***<u>Supplementary Figure 9:</u> ERPs from sites in the Inferior Occipital Cortex (IOC) responding to social cues.*** *The patient, electrode and site number, as well as the anatomical localization and the MNI coordinates are indicated for each ERP. Horizontal green bars indicate the time window where the conditions significantly differ. Black diamonds near the x-axis of the plots indicate polarity inversions in adjacent sites. Legend: IOS: Inferior Occipital Sulcus, LOS: Lateral Occipital Sulcus, MOG: Mid-Occipital Gyrus, tpCollS: Transverse Posterior Collateral Sulcus. *: p<0.05, **: p<0.01, ***: p<0.005, Monte-Carlo p-values.*

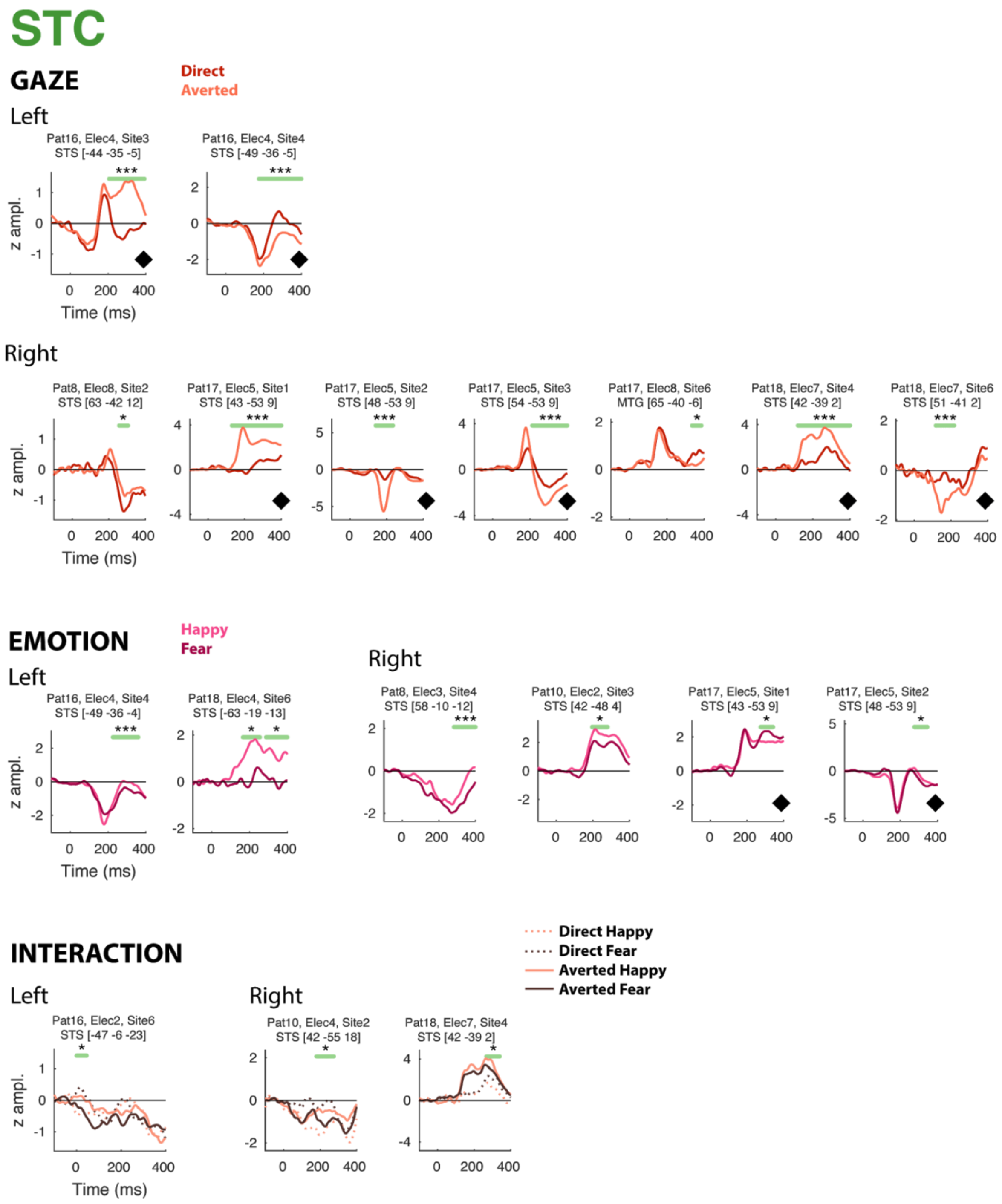


**<u>Supplementary Figure 10:</u> *ERPs from sites in the Superior Temporal Cortex region (STC) responding to social cues.*** *The patient, electrode and site number, as well as the anatomical localization and the MNI coordinates are indicated for each ERP. Horizontal green bars indicate the time window where the conditions significantly differ. Black diamonds near x-axis indicate polarity inversions in adjacent sites. MTG: Mid-Temporal Gyrus, STS: Superior Temporal Sulcus. *: p<0.05, **: p<0.01, ***: p<0.005, Monte-Carlo p-values.*

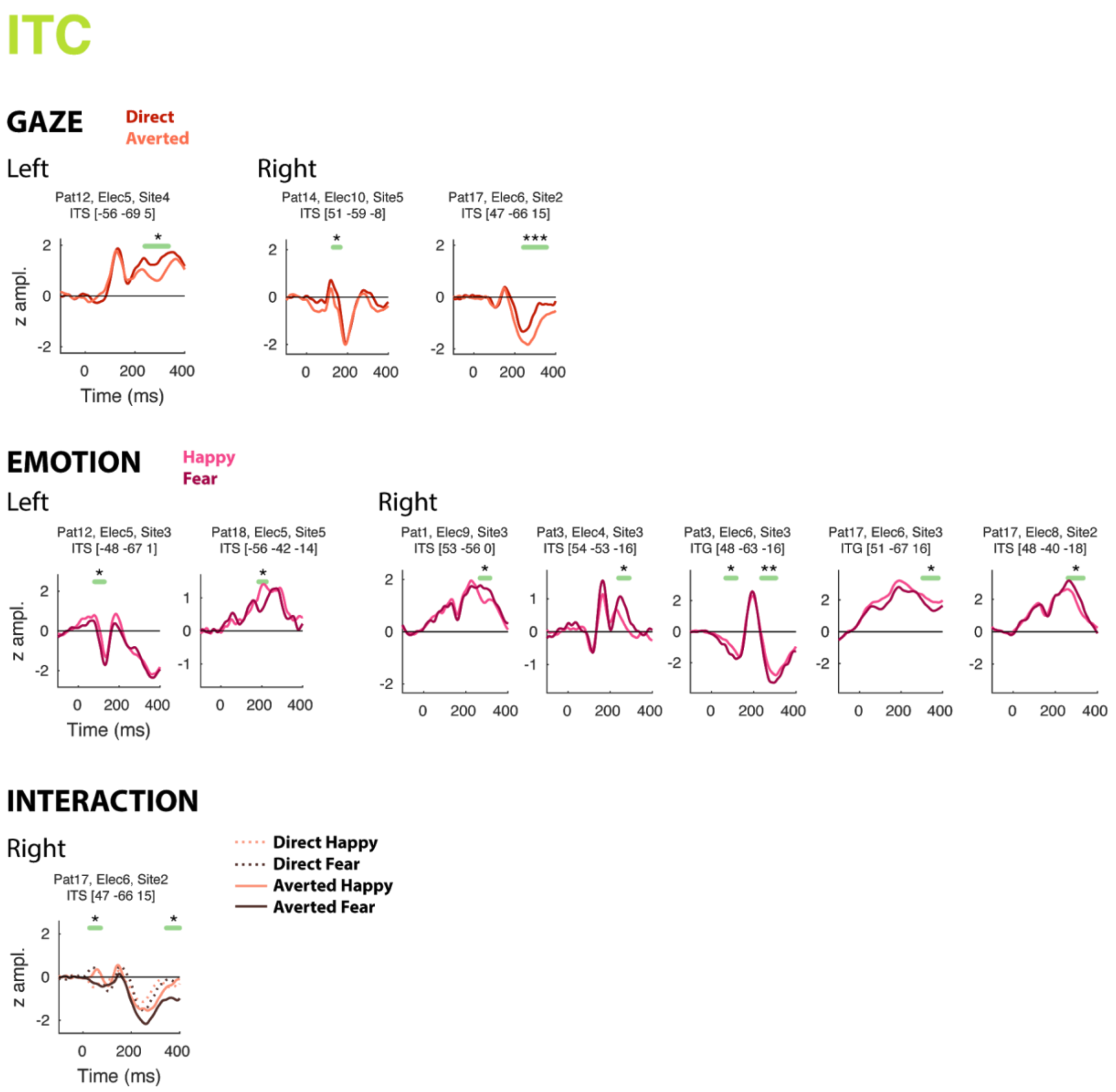


**<u>Supplementary Figure 11:</u> ERPs from sites in the Inferior Temporal Cortex region (ITC) responding to social cues.** *The patient, electrode and site number, as well as the anatomical localization and the MNI coordinates are indicated for each ERP. Horizontal green bars indicate the time window where the conditions significantly differ. Legend: ITG: Inferior Temporal Gyrus, ITS: Inferior Temporal Cortex. *: p<0.05, **: p<0.01, ***: p<0.005, Monte-Carlo p-values.*

*S2.10. Additional check regarding stimulus attributes and trial numbers*

It should be noted that at the level of the physical stimulus, the gaze changes were very small compared to the overall change that the emotion produced within the whole face (see Supplementary Fig. 2 of (Huijgen et al. 2015)). We made sure that our gaze effects were not confounded by emotion, which could potentially be the case if the final number of trials in the 2-by-2 design was imbalanced. None of the patients showing gaze effects had an imbalanced number of trials per condition (Fisher's exact tests for each patient on the number of trials for each condition taken 2 by 2: all p>0.45, uncorrected for multiple comparisons across patients tested), thereby ruling out this potential confound.

*S2.11. Effects of the direction of gaze change*

We found three sites sensitive to the direction of gaze change (see Supplementary Fig.12 below). Two of these sites were located in the right STS (Patient 17) and presented larger ERPs for contralateral gaze direction (cluster-based permutation t-test; Electrode 5, site 1: cluster sum of t=55.1, p=0.017; Electrode 5 site 3: cluster sum of t=30.3, p=0.043; uncorrected for multiple comparisons over sites tested). The third site was located in the left ITS (Patient 12) and presented an additional early negativity for ipsilateral gaze direction (cluster sum of t=-75.7, p=0.0068).

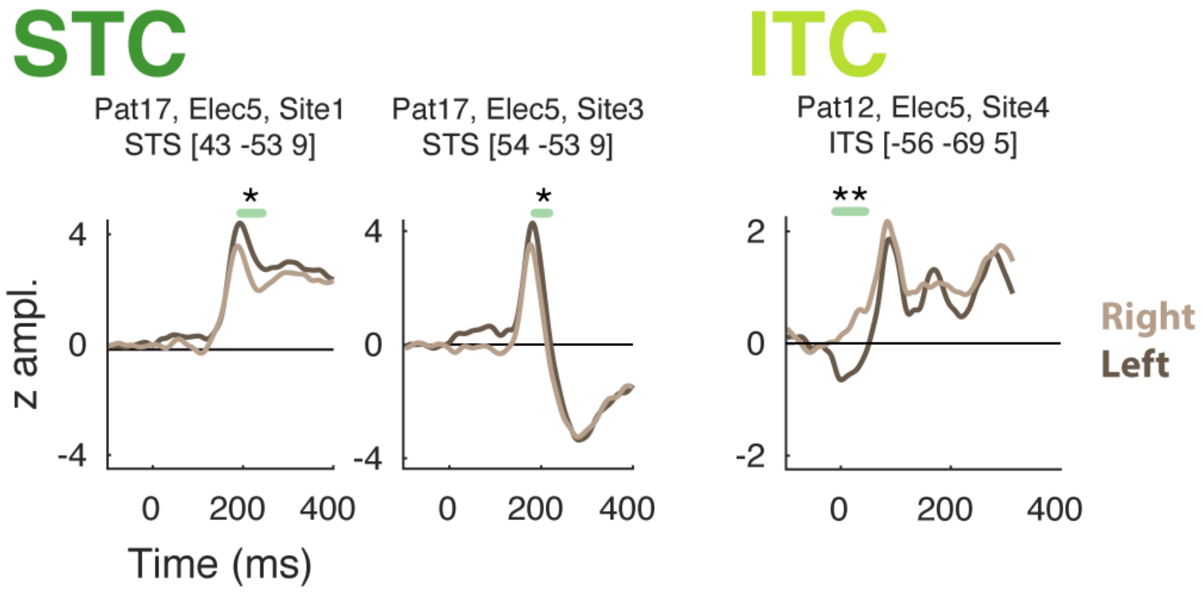

***Supplementary Figure 12:*** ***left vs right gaze effects.*** *The patient, electrode, site number, anatomical label and the MNI coordinates are indicated for each ERP, whose amplitudes are depicted in terms of z-scores. Horizontal green bars indicate the time window where the conditions significantly differ. Legend: ITC: Inferior Temporal Cortex, ITS: Inferior Temporal Sulcus, STC: Superior Temporal Cortex, STS: Superior Temporal Sulcus. *: p<0.05, **: p<0.01, Monte-Carlo p-values.*
